# Deciphering Functional Heterogeneity of Cancer-Associated Fibroblasts Across Molecular Subtypes of Breast Cancer

**DOI:** 10.1101/2025.01.05.631269

**Authors:** Dharambir Kashyap

## Abstract

Breast cancer heterogeneity has been well understood based solely on tumor components. However, the tumor microenvironment, which plays a crucial role in cancer by regulating tumor cells’ differentiation, maturation, and malignant potential, remains under-explored. This study conducted high-throughput RNA-Seq analysis on fresh breast cancer tissues from each molecular subtype and a pure population of cancer-associated fibroblasts (CAFs) isolated from cancer tissue and adjacent normal parts. Based on their functions, we identified three populations of CAFs: a) immune response-related, b) ECM remodeling, and c) calcium/protein binding. Validation of differentially expressed genes among these CAF populations revealed subtype-specific correlations. For instance, CAFs expressing immune response-related genes were significantly enriched in Luminal A/B and TNBC (log +1-fold; p =0.00047), compared to Her-2 positive (log -2.3-fold; p =0.0026). ECM remodeling genes exhibited greater expression in Her-2 positive and TNBC (log +4 to +11-fold; p =7.87E-08) when compared to Luminal subtypes (log +1.5-fold to +6-fold; p =8.66E-05). Calcium binding genes displayed overall similar expression across breast cancer subtypes. Thus, this study identified and underscored the functional heterogeneity of CAFs within the tumor microenvironment across the molecular subtypes of breast cancer.

## 1. Introduction

Breast cancer is a heterogeneous disease at the cellular and molecular levels, as shown by gene expression profiling, which defines its various molecular subtypes [1–3]. This heterogeneity has been analyzed based on the tumor component of breast cancer [4]. However, the tumor microenvironment (TME), a crucial part of cancer that regulates processes such as differentiation, maturation, and the malignant potential of tumor cells, remains underexplored [5–7]. The complexity of TME components and their interactions with surrounding neoplastic cells create a cascade of changes in multiple pathways that may have lethal repercussions [8, 9].

In the last decade, the focus has shifted from tumor cells to TME to address their complexity and elucidate their role in breast carcinoma progression and response to therapy [10, 11]. TME itself is a highly complex system mainly composed of tumor cells, immune Infiltrating cells, such as macrophages, dendritic cells, lymphocytes, cancer-associated fibroblasts (CAFs), endothelial cells, and adipocytes, as well as the extracellular matrix (ECM) and various signaling molecules [5, 6]. CAFs are the most abundant and essential stromal components in the tumor microenvironment (TME). Previous findings have shown that CAFs play a role in multiple stages of tumor development through diverse pathways [11–13]. The bidirectional crosstalk between CAFs and tumor cells, along with other cell types, is mediated by CAF-derived cytokines, chemokines, growth factors, and exosomes within the TME, facilitating tumor proliferation and inducing immune evasion by cancer cells [14–16]. CAFs can also degrade ECM by releasing matrix metalloproteinases (MMPs) while synthesizing new matrix proteins to provide structural support for tumor invasion and angiogenesis [17]. However, more specific roles and detailed mechanisms of CAFs in cancer pathogenesis and progression need further exploration. These crucial cells are highly heterogeneous in their origin, function, and phenotype, possessing various endogenous and surface markers, including vimentin, NG2, platelet-derived growth factor receptor (PDGFR)α/β, α-smooth muscle actin (α-SMA), fibroblast activation protein (FAP), podoplanin (PDPN), and fibroblast specific protein 1 (FSP1) [18]. These markers are vital for the pro-tumorigenic functions of these cells [18]. These biomarkers are not explicitly linked to a specific CAF or cancer type [18]. Hence, a heterogeneous population of CAFs exists, characterized by various phenotypes and distinct functions across many cancer types. For example, several studies have previously identified functionally distinct subtypes of CAFs in different human cancers, such as pancreatic and breast cancer. Ohlund et al. (19) discovered two subsets of CAFs, myofibroblastic CAFs and inflammatory CAFs, within pancreatic cancer (19). Another study identified a new CAF subpopulation that expresses MHC class II and CD74, termed antigen-presenting CAFs [20]. Similarly, distinct CAF subpopulations have been reported in breast cancer. One study classified four CAF subsets (S1-S4) based on the expression of CD29, FAP, α-SMA, PDGFRβ, FSP1, and caveolin-1. Bartoschek et al. [21] also identified four subpopulations of CAFs (vCAFs, mCAFs, cCAFs, and dCAFs) according to their distinct cellular sources using single-cell RNA sequencing [21]. As a result, a deeper understanding of the heterogeneity of CAFs in breast cancer is necessary. Therefore, we aimed to decipher the heterogeneity of CAFs across molecular subtypes of breast cancer by analyzing the whole transcriptome of tumor tissue and the Ampli-Seq transcriptome of pure CAFs.

## 2. Materials and Methods

### Sample collection

The study received approval from the Institutional Ethical Committee. Informed consent was obtained from each patient participating in the study. Breast cancer tissue from each subtype was collected from treatment-naive patients undergoing mastectomy or lumpectomy procedures. Each case was processed into formalin-fixed paraffin-embedded sections and surrogated for molecular subtyping using immunohistochemistry (IHC) for estrogen receptor, progesterone receptor, Her-2 neu, and Ki67. Based on the IHC pattern, cases were categorized as Luminal A, Luminal B, Her-2 positive, and Triple Negative Breast Cancer (TNBC).

### Isolation of CAF

CAFs were isolated from fresh breast cancer tissue and normal breast tissue (taken 5 cm away from the cancer tissue) using an enzymatic method. Briefly, 100 mg of tissue was chopped in an enzyme mix solution containing 200 units/mL of collagenase type I (cat. 17018029, Gibco) and 500 units/mL of hyaluronidase (cat. H3506, Sigma Aldrich) in growth media, incubating at 37°C for 1.5 hours. After digestion, the tissue lysate was filtered through a 40 µm cell strainer and centrifuged at 400 X g for 5 minutes. The obtained cell pellet was seeded in a 6-well culture plate containing growth media with Dulbecco’s Modified Eagle Medium/F12 (cat. 11320033, Gibco), 10% FBS (cat. 10082147, Gibco), and 1% antibiotic and antimycotic solution (Cat. A002, Hi-Media), and maintained at 37°C with a constant supply of 5% CO2. After 2 hours, the attached cells were gently washed with PBS and nourished with fresh growth media. The growth media was changed alternately, and cells were passaged every 3-4 days up to 7 passages. (Supplementary Figure 1 shows the isolated CAFs and normal breast fibroblasts).

### RNA Isolation

Total RNA was isolated using mirVana™ RNA isolation kit (Cat. AM1560, Ambion) from fresh frozen breast cancer tissue, primary cultured CAFs, and normal fibroblasts. ReliaPrep™ FFPE Total RNA miniprep system kit (Cat No. Z1002, Promega) was used for total RNA isolation from formalin-fixed and paraffin-embedded (FFPE) breast cancer samples.

### Transcriptomic Profiling

Whole transcriptome RNA-Seq was conducted on four fresh breast cancer tissues, with one case from each subtype. Additionally, AmpliSeq transcriptome analysis was performed on pure cultures of CAFs and normal breast fibroblasts isolated from each breast cancer subtype and normal breast tissue (Luminal A 1, Luminal B 2, Her-2+ 2, TNBC 2, normal breast 1). The AmpliSeq transcriptome utilized a human gene expression panel (Cat A26325, Thermo Fisher Scientific). For both RNA-Seq runs (whole transcriptome and AmpliSeq transcriptome), the Ion Total RNA-Seq Kit v2.0 (Cat No. 4475936, Thermo Fisher Scientific) was used for library preparation, and RNA sequencing was conducted on Ion S5 systems from Thermo Fisher Scientific, following the manufacturer’s instructions.

### Data analysis

Raw data from both RNA-Seq runs was converted to FASTQ format and utilized by the AmpliSeqRNA plugin for further analysis. Sequencing reads were aligned to the reference genome, human GRCh38, for annotation using TMAP software with default parameters. Differential expression analysis between sample groups was conducted with the DESeq2 tool from Bioconductor in the R package. Genes with ≥10 read counts, p-value ≤0.05, and log2-fold change ≥2 were included for functional gene enrichment analysis. Online tools, including the Database for Annotation, Visualization and Integrated Discovery (DAVID) Bioinformatics Resources 6.8, PANTHER 16.0 GSEA, and FunRich (Functional Enrichment Analysis Tool), were used for gene functional enrichment analysis.

### Gene selection for validation

Differentially expressed stromal genes in various molecular breast cancer subtypes were selected based on maximum expression fold, p-adjusted value >0.005, and functional significance (Table 1).

**Table 1.**
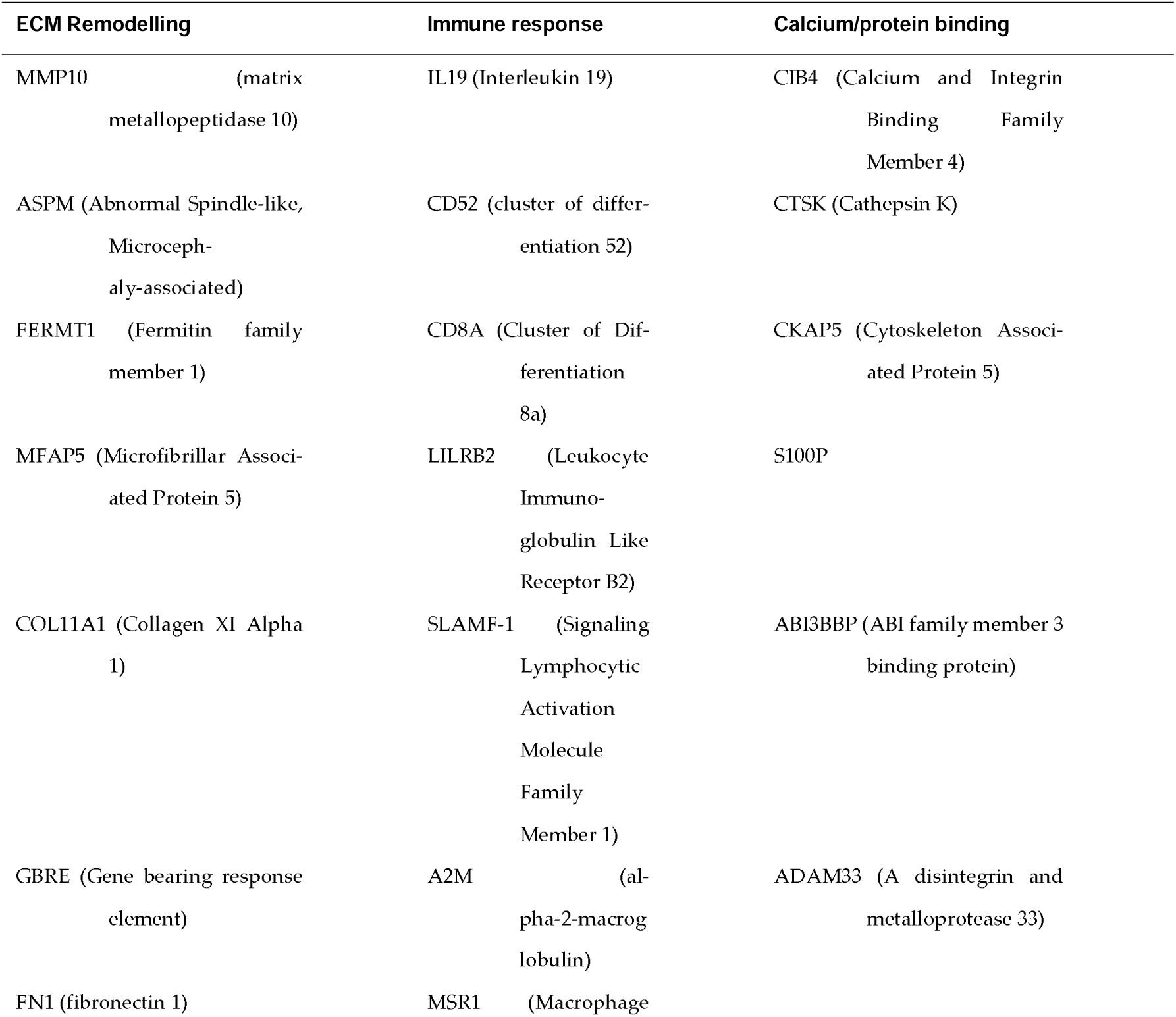
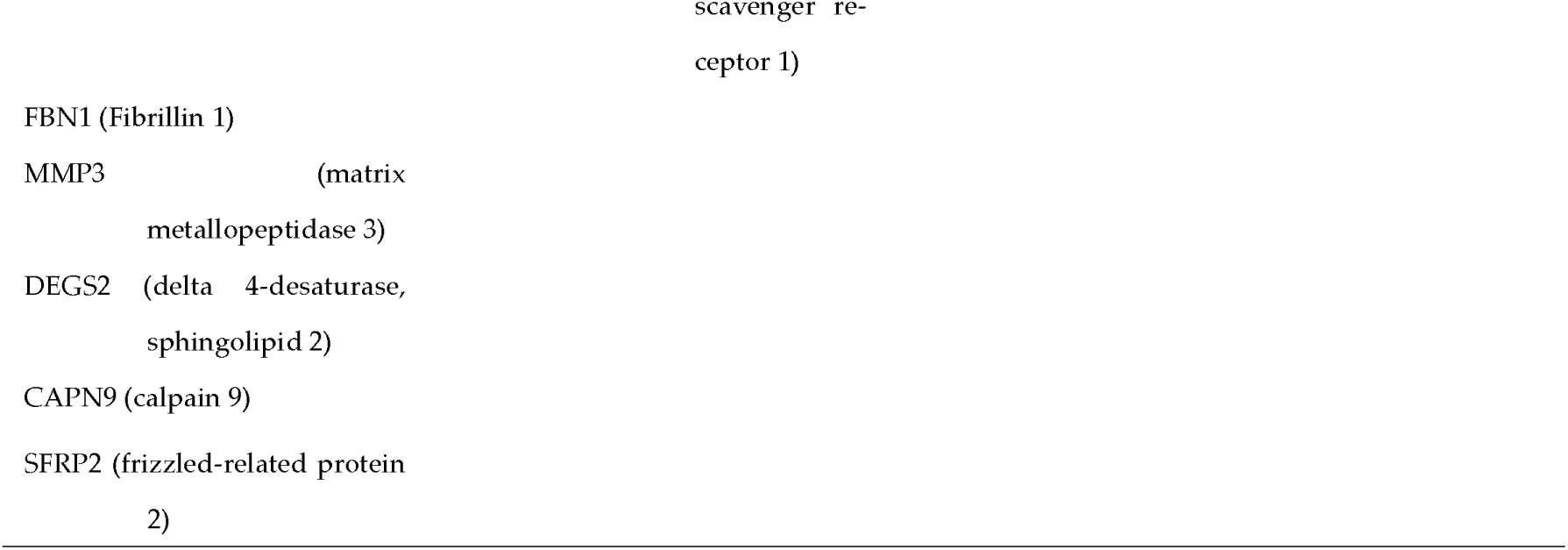
Grouping of the genes based on their biological functions

Validation of shortlisted genes: Shortlisted genes were validated on 83 FFPE breast cancer samples (Luminal A-25, Luminal B-18, Her-2 positive-18, TNBC-22). To achieve this, the corresponding H&E slide of each selected tissue block was microscopically analyzed to ensure adequate stroma. Total RNA (1 µg) was isolated from the selected stromal part of the tumor block and utilized for cDNA synthesis. Real-time qPCR reactions were conducted using the Brilliant III Ultra-Fast SYBR Green QPCR Master Mix-Agilent on the CFX96 Touch Real-Time PCR Detection System (Bio-Rad). Three housekeeping genes (ß-actin, RPL-4, and RPS-1) served as references for normalization. All reactions were performed in triplicate and calculated using the 2ΔΔCt formula.

## 3. Results

### 3.1.1. Whole transcriptome (WT)

Approximately 16,000 genes were mapped in each molecular subtype of breast cancer. For subsequent analysis, we focused only on statistically significant genes (p≥ 0.05). Venn diagrams (Fig. 1a and 1b) display several significantly differentially expressed genes (DEGs) in each subtype and within subtypes. The DESeq2 tool used for differential gene expression analysis among the breast cancer subtypes identified a distinct set of CAFs-derived genes significantly associated with the molecular subtypes. The heatmap generated with the top 30 DEGs illustrated the genes related to the subtypes (Fig. 2). Since WT was conducted on bulk tumor tissues, only genes belonging to the stroma were included for final analysis. Functional gene set enrichment analysis associated DEGs with various cancer-promoting molecular functions and cancer signaling pathways. Most of the significantly expressed genes (adjusted p-value≥ 0.05, FDR p ≥ 0.05) were associated with extracellular matrix (ECM) remodeling components, ECM adhesion components, and immune modulators, such as protease enzymes. In the Luminal A subtype, significantly upregulated genes (log +3.1 to +5.4 -fold; adjusted p = 0.0022) were involved in cell differentiation, regulation of glucose import/metabolism, pH regulation, protein kinase A and C signaling, and G-coupled receptor signaling pathways. Additionally, significantly downregulated genes (log -2.4 to -2.0-fold, p= 0.027) were found to be involved in the ubiquitination system, RNA binding proteins, glucosamine enzyme activities, and translation elongation factors, among others (Fig. 3A). In the Luminal B subtype, significantly upregulated genes (log +6.9 to +2.2-fold; adjusted p = 0.00047) were enriched in programmed necrotic cell death pathways, immune modulation functions, and drug response signaling pathways. Conversely, significantly downregulated genes (log -3.8 to -2.6 -fold, p = 0.00026) primarily featured RNA molecule binding functions (Fig. 3B). The stroma of Her-2 positive tumors exhibited significantly upregulated genes (log +3.3 to +10-fold; adjusted p = 1.22E-13) with roles in ECM organization, cell-matrix adhesion, vasoconstriction, blood coagulation factors, and regulators of ERK1/2 signaling pathway activities. Downregulated genes (log -6.7 to -3.8-fold; adjusted p = 7.897E-08) included sphingosine hydroxylase, sphingolipid desaturase, glutaminase, and acetylesterase, among others (Fig. 3C). TNBC subtypes revealed significantly upregulated genes (log +2.6 to +9.4-fold; adjusted p= 3.96E-10) that were involved in fibroblast growth factor receptor signaling pathways, transmembrane transporters, apoptosis inhibitor activity, positive regulation of proliferation, and oxidation-reduction molecular processes. Downregulated genes (log -11 to -5.6-fold; adjusted p= 5.39E-09) mostly functioned as cell-cell adhesion molecules, B-cell receptor signaling pathways, phagocytosis processes, innate immune response, and G-protein coupled receptor signaling pathways, among others (Fig. 3D).

**Figure-1:**
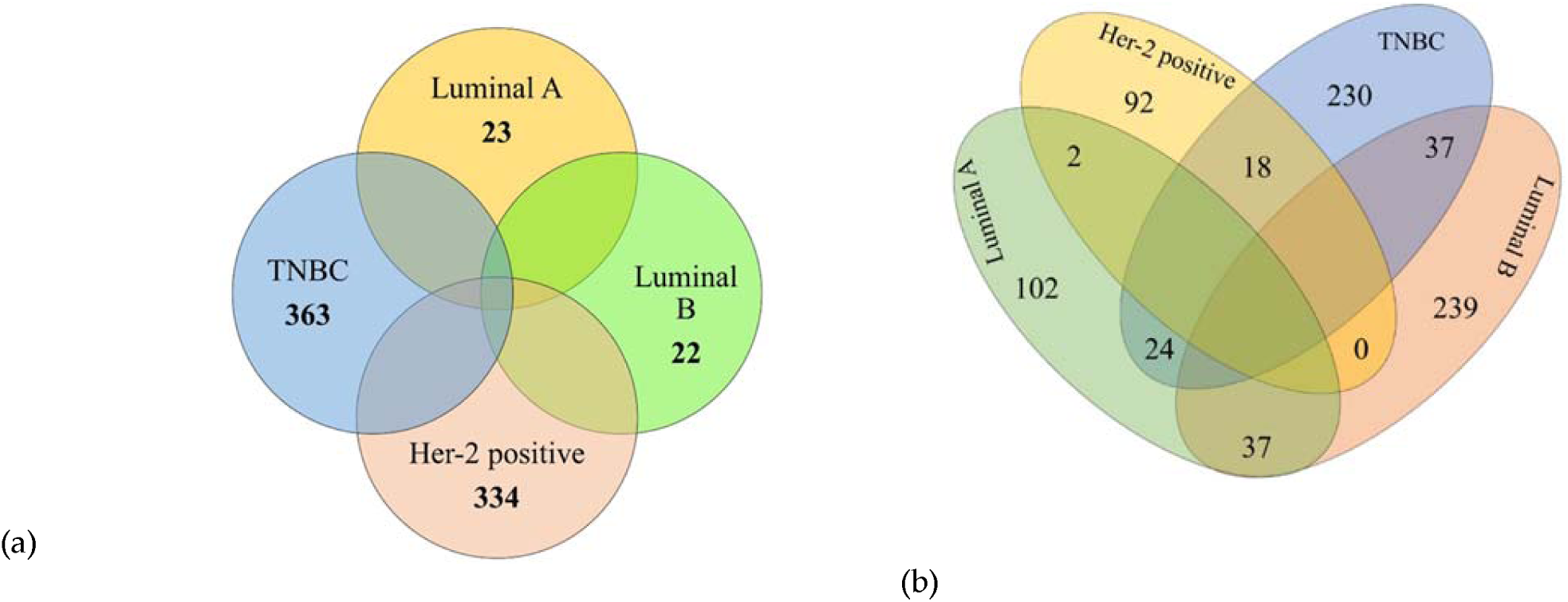
**(a)** Venn diagram of upregulated genes from Whole Transcriptome; (b) Venn diagram of downregulated genes from Whole Transcriptome

**Figure 2:**
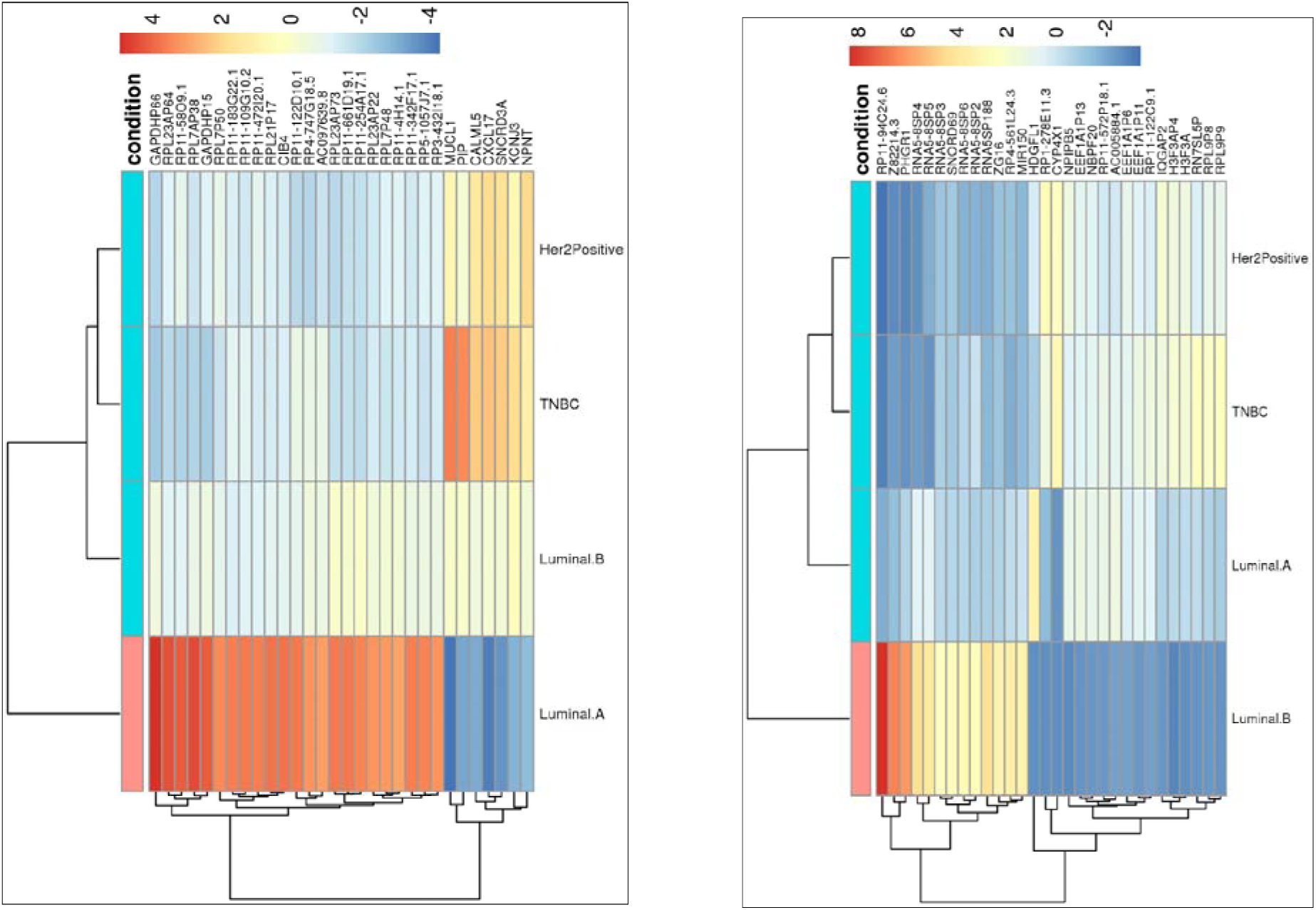
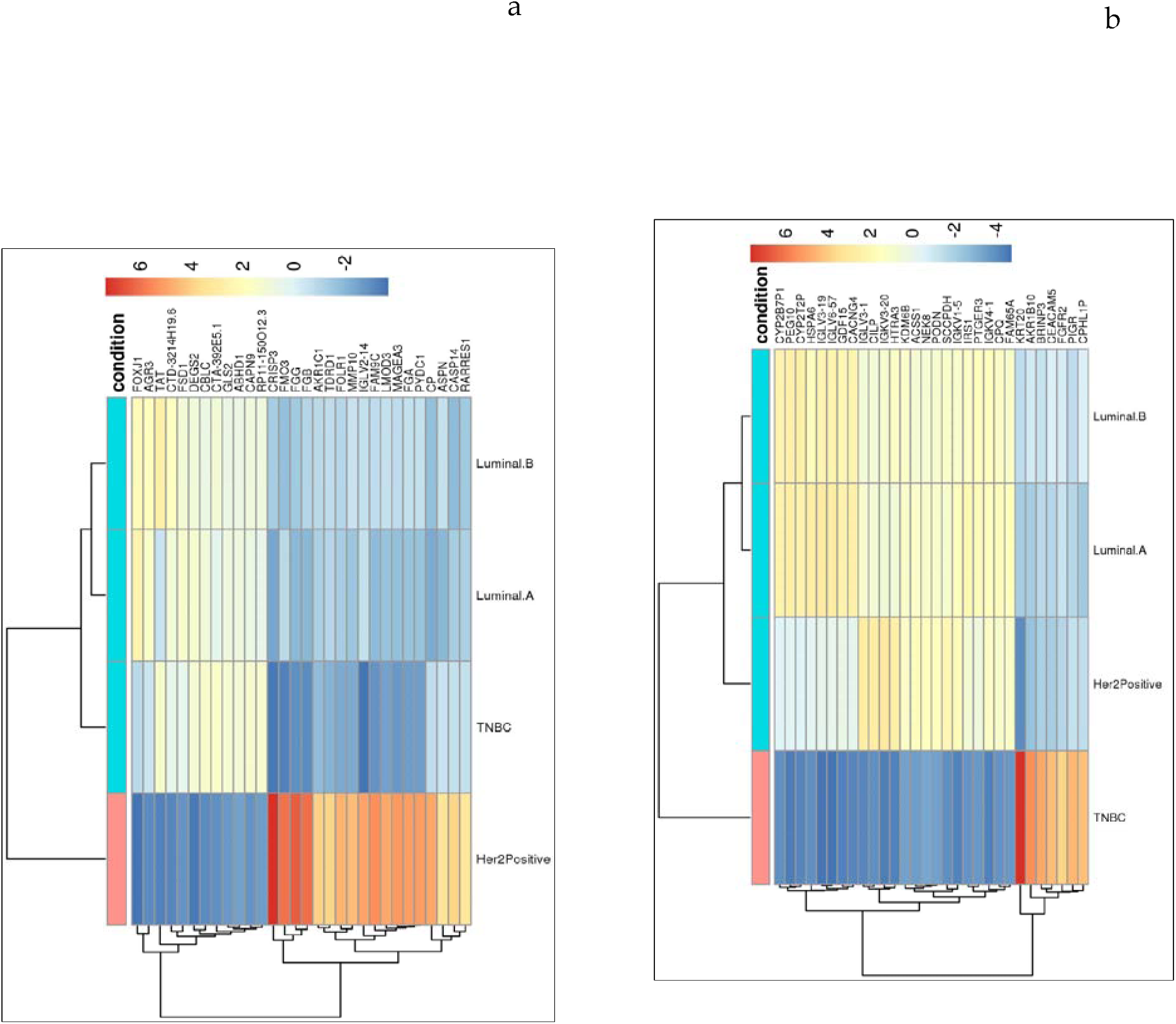
Heatmaps generated with top DEGs from Whole transcriptome data. (a-d) Heatmap showing the DEGs between the breast cancer subtypes.

**Figure-3A:**
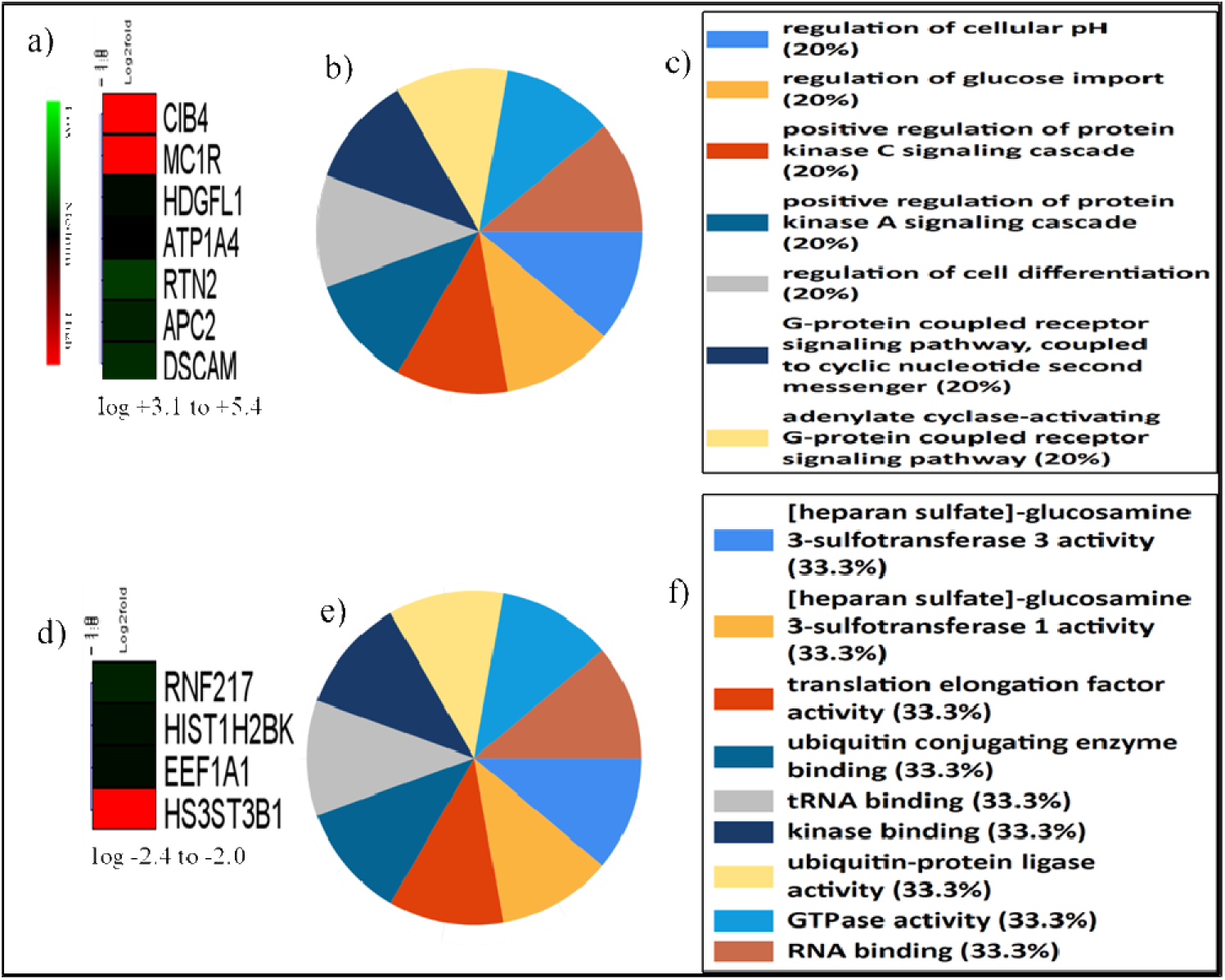
Differentially expressed genes from the whole transcriptome in the Luminal A subtype: a) Heatmap of upregulated DEGs, b) Pie charts displaying the functional distribution of upregulated DEGs, c) Various functions of the upregulated DEGs, with most related to pH regulation and G-protein coupled receptor signaling; d) Heatmap of downregulated DEGs, e) Pie charts displaying the functional distribution of downregulated DEGs, f) Various functions of the downregulated DEGs, including heparan sulfate-glucosamine and ubiquitin-conjugating enzyme.

**Figure 3B:**
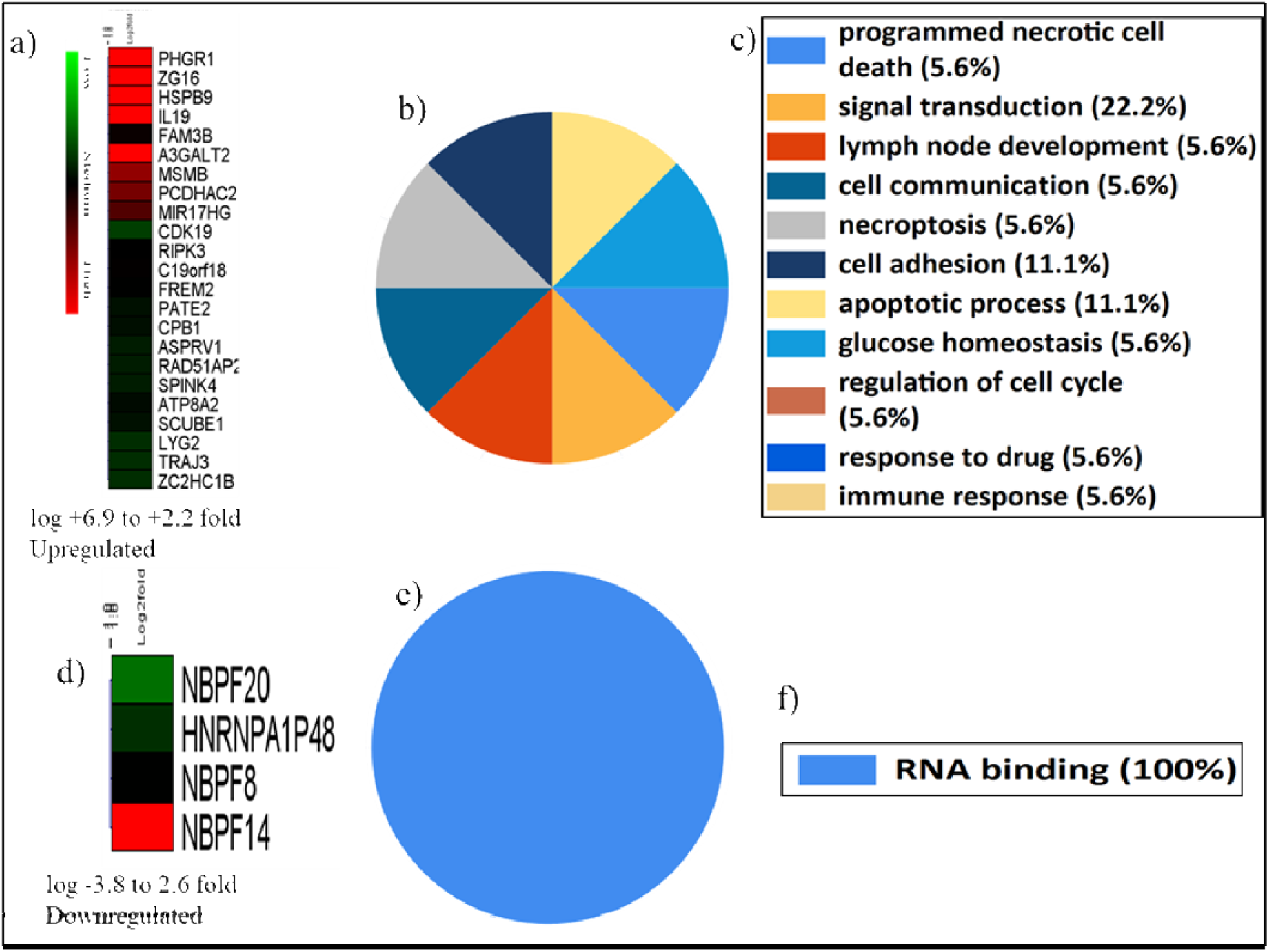
Differentially expressed genes from Whole transcriptome in Luminal B subtype, a) Heatmap of upregulated DEGs, b) Pie charts showing the distribution of functions of upregulated DEGs, c) Different functions of the upregulated DEGs. Most of the functions are related to cell adhesion, immune response, and response to drugs, Heatmap of downregulated DEGs, e) Pie charts showing the distribution of functions of downregulated DEGs, f) Different functions of the downregulated DEGs.

**Figure-3C:**
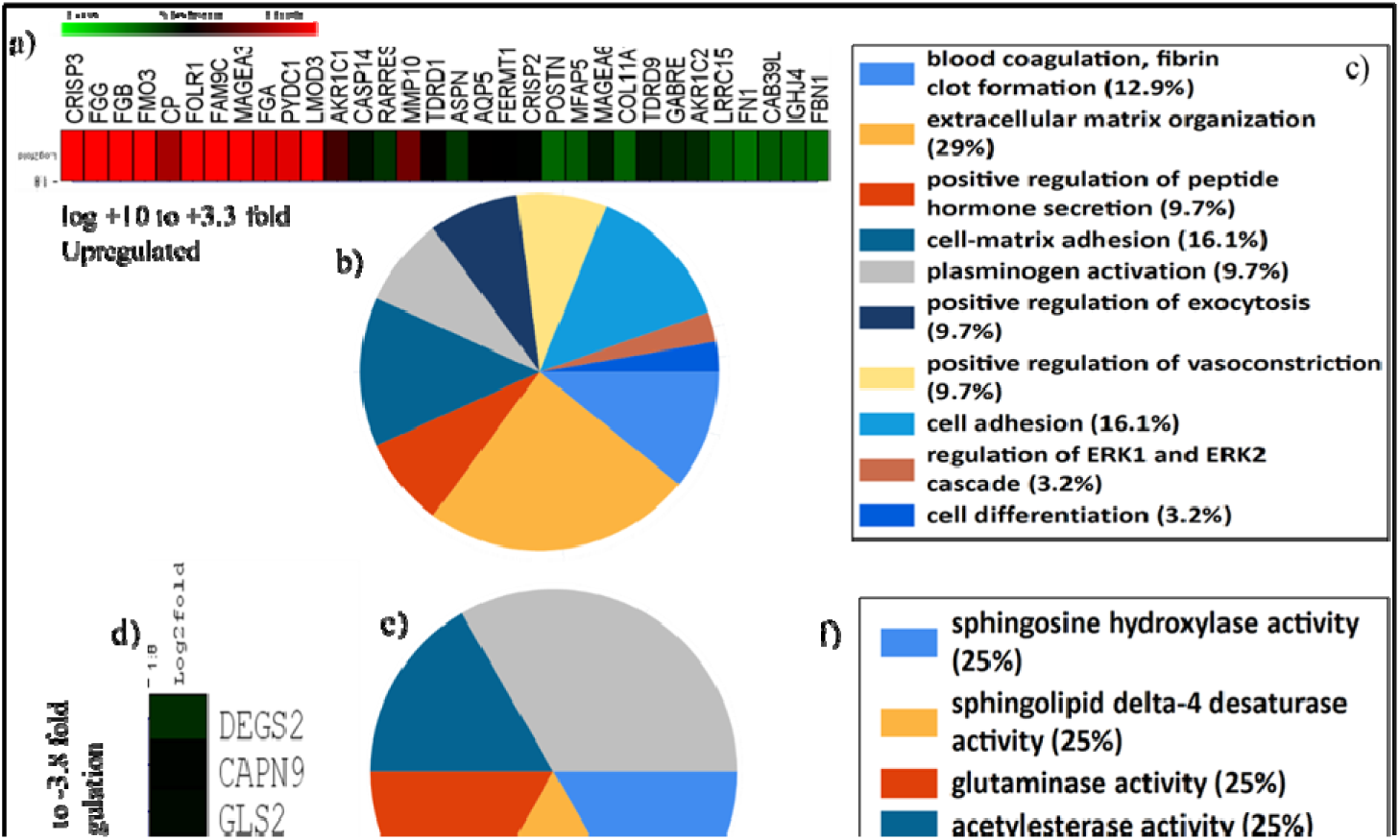
Differentially expressed genes from the whole transcriptome in the Her-2 positive subtype: a) Heatmap of upregulated DEGs, b) Pie charts showing the distribution of functions of upregulated DEGs, c) Different functions of the upregulated DEGs; most functions relate to ECM regulation, cell-matrix adhesion, and plasminogen activation, d) Heatmap of downregulated DEGs, e) Pie charts showing the distribution of functions of downregulated DEGs, f) Different functions of the downregulated DEGs, including cell sphingosine hydroxylase activity and sphingolipid delta-4 desaturase.

**Figure-3D:**
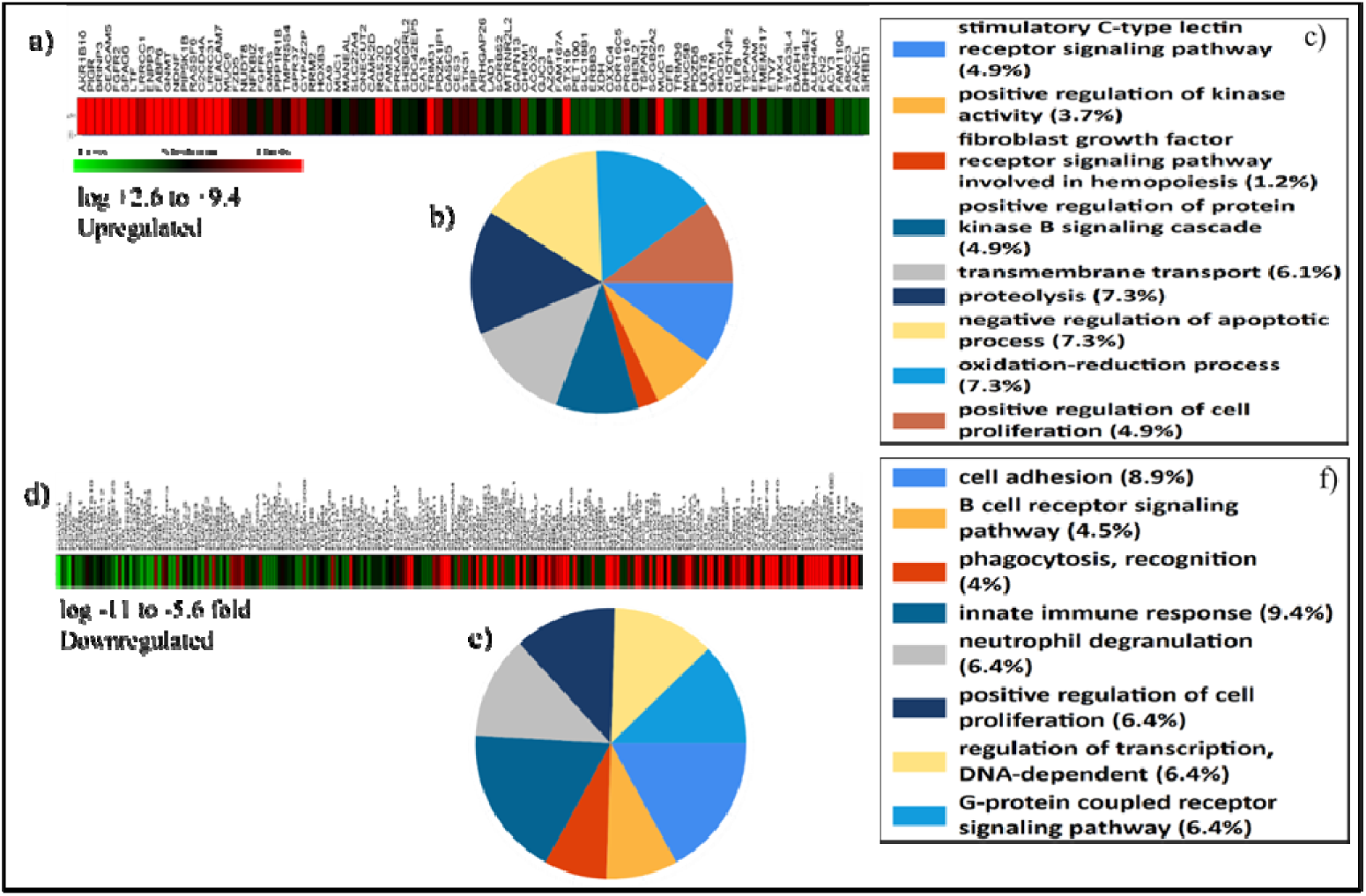
Differentially expressed genes from the whole transcriptome in TNBC subtype: a) Heatmap displaying upregulated DEGs, b) Pie charts illustrating the distribution of functions of upregulated DEGs, c) Various functions of the upregulated DEGs; most functions are related to fibroblast growth factor receptors, transmembrane transport, and oxidation-reduction, d) Heatmap showing downregulated DEGs, e) Pie charts depicting the distribution of functions of downregulated DEGs, f) Various functions of the downregulated DEGs; which include cell adhesion, innate immune response, and neutrophil degradation.

### 3.1.2. AmpliSeq transcriptome

Approximately 14,000 genes were mapped in different breast cancer subtypes. We focused on statistically significant genes (p≥ 0.05) for subsequent analysis. Venn diagrams (Fig. 4a and 4b) illustrate the number of statistically significant DEGs in each subtype and the overall count within subtypes. DEG analysis (using the DESeq2 tool) was conducted between breast cancer samples and normal breast tissue, as well as among breast cancer subtypes. This analysis also identified distinct CAFs-derived genes significantly associated with molecular subtypes. The heatmap generated with the top 30 DEGs displayed the genes associated with each subtype (Fig. 5). In the functional gene set enrichment analysis, the Luminal A subtype exhibited that their upregulated genes (log +3.1 to +2.4 fold; p=0.015) had functions related to metal and protein binding activity, methyltransferase activity, MAP kinase phosphatase activity, transcription factor binding, metal binding, binding to collagen or other ECM proteins, and the activation of autophagosome proteins; their downregulated genes (log -2.4 fold; p=0.072) were enriched in lamin binding, phospholipase C activity, and calmodulin binding activities (Fig. 6A). Upregulated DEGs in the Luminal B subtype (log 2.0 to +2.1-fold; p adjusted=0.0005) were enriched in collagen, heparin, ECM protein binding activities, receptor tyrosine kinase, fatty acid metabolism, and ephrin receptor-mediated signaling (Fig. 6B). Downregulated genes in Luminal B were statistically non-significant and thus excluded from functional analysis. Additionally, significantly upregulated genes (log +4.0 to +6.0-fold; p=0.031) in Her-2 positive tumors were noted for functions related to G protein-coupled receptor activity, olfactory receptors, angiotensin receptors, and leukotriene receptor-mediated activities, while downregulated genes (log -3.7 to -5.1-fold; p=0.018) exhibited functions such as ECM structural components, collagen binding components, ribonucleoprotein complex binding proteins, and binding proteins for collagen fibers and microtubules (Fig. 6C). Lastly, TNBC tumors showed a significant upregulation of genes (log +5.7 to +6.4-fold; p adjusted=0.0062) enriched in ECM-supporting components (fibronectin) and proteins that regulate Wnt, calcium, and low-density lipoproteins, among others. Other DEGs in TNBC were involved in cargo receptor and RAGE receptor-regulated processes, as well as extracellular binding signaling pathways (Fig. 6D). Downregulated genes in TNBC were statistically non-significant and thus excluded from functional analysis analysis.

**Figure-4:**
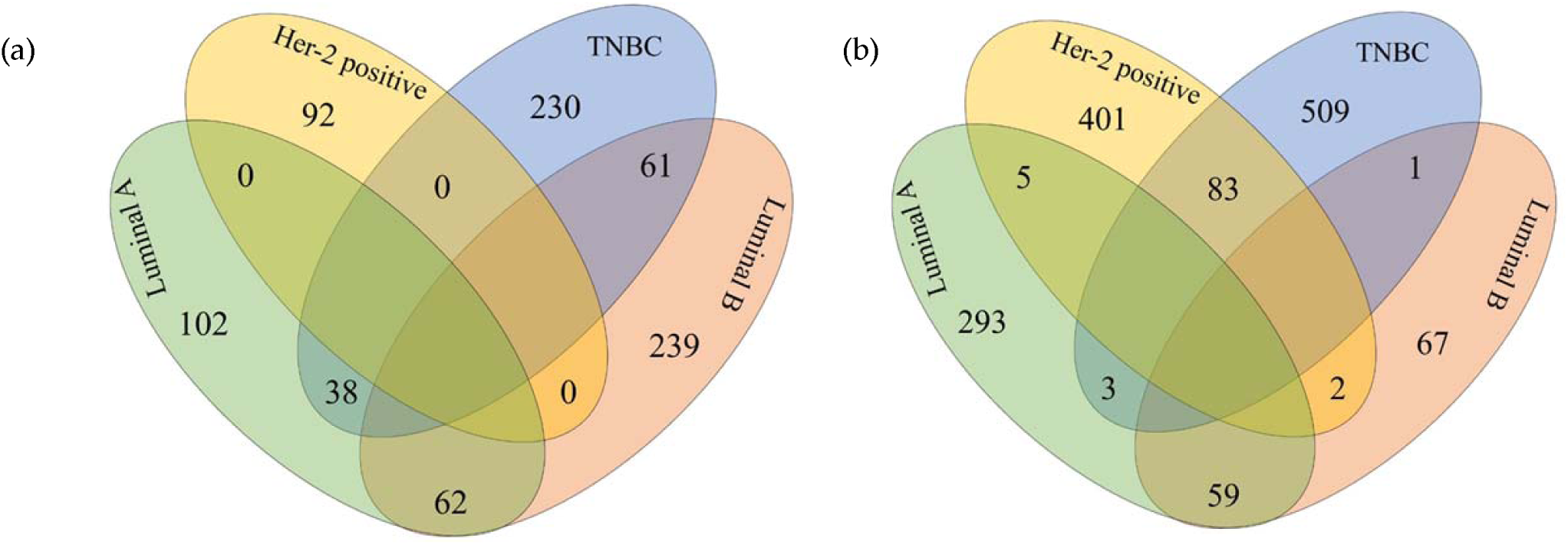
(a) Venn diagram of upregulated genes from Ampliseq (Human ampliseq targeted panel); (b) Venn diagram of downregulated genes from Ampliseq (Human ampliseq targeted panel)

**Figure-5:**
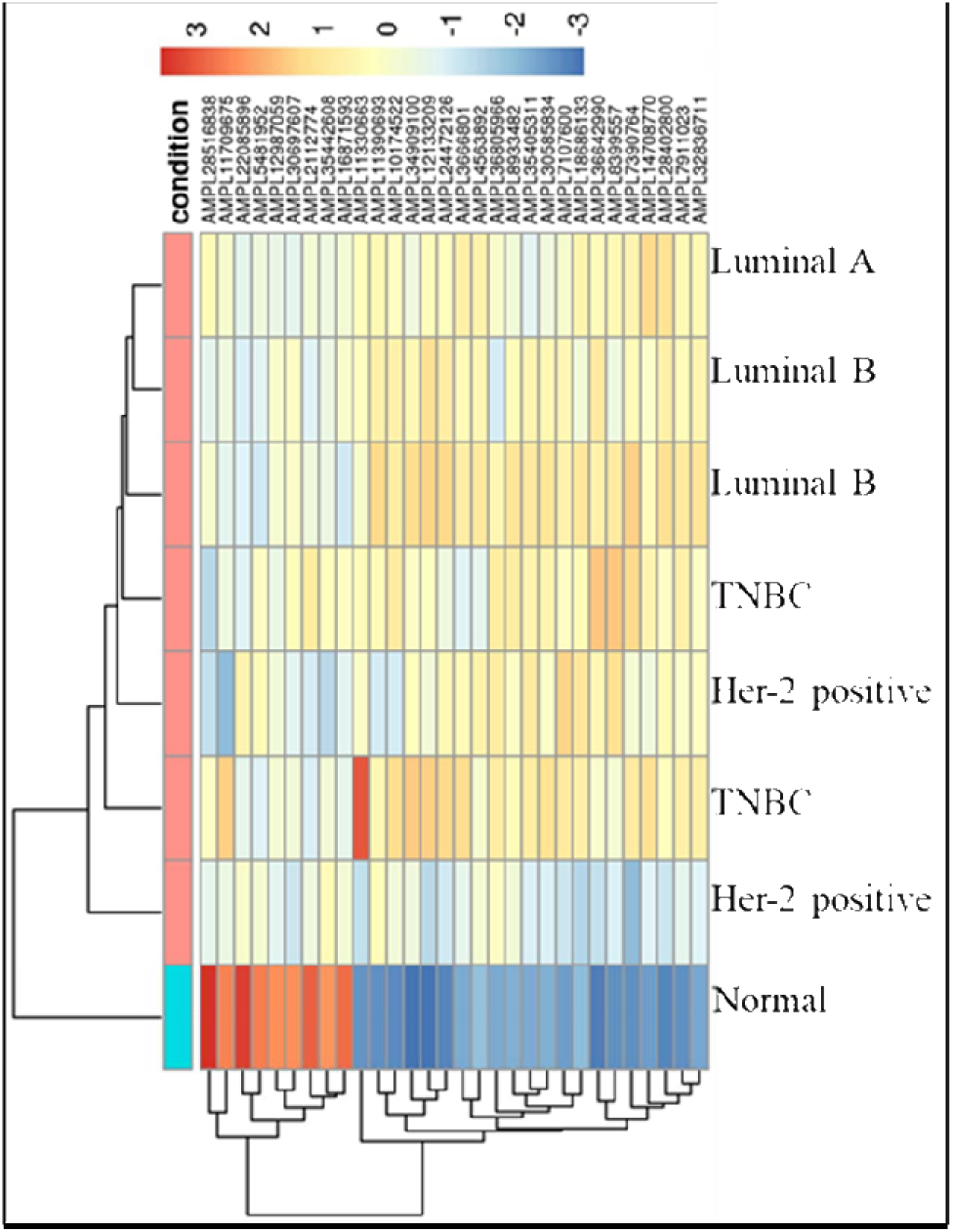
Heatmaps generated with top 30 DEGs from AmpliSeq transcriptome data from the breast cancer subtypes.

**Figure-6A:**
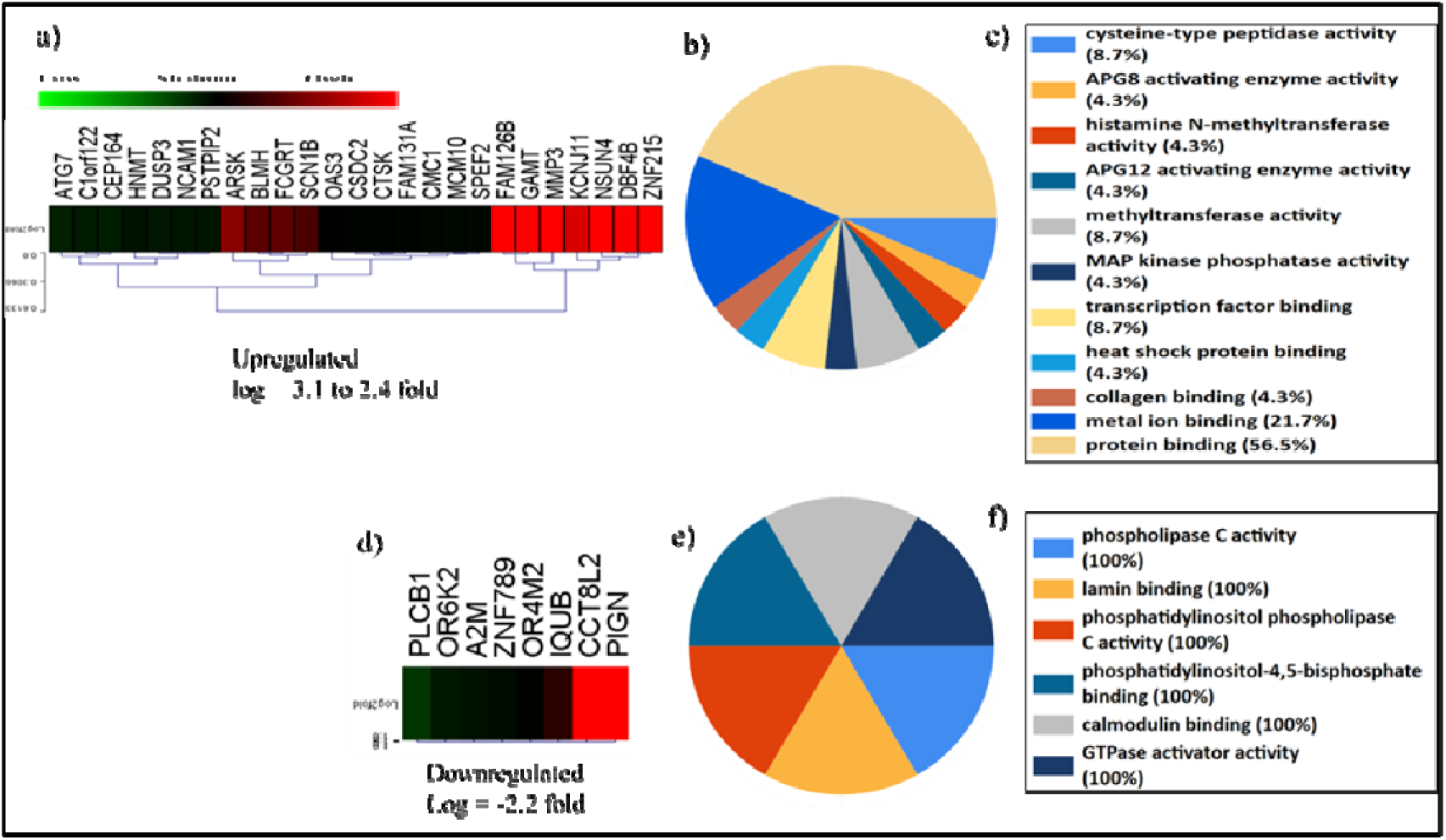
Differentially expressed genes from the AmpliSeq transcriptome in the Luminal A subtype. a) Heatmap of upregulated DEGs, b) Pie charts illustrating the distribution of functions of upregulated DEGs, c) Various functions of the upregulated DEGs. Most genes exhibited protein binding, MAP kinase phosphatase activity, and collagen binding functions. Additional functions included methyltransferase activity and histamine N-methyltransferase activity, d) Heatmap of downregulated DEGs, e) Pie charts illustrating the distribution of functions of downregulated DEGs, f) Various functions of downregulated DEGs, including calcium binding and lamin binding function.

**Figure-6B:**
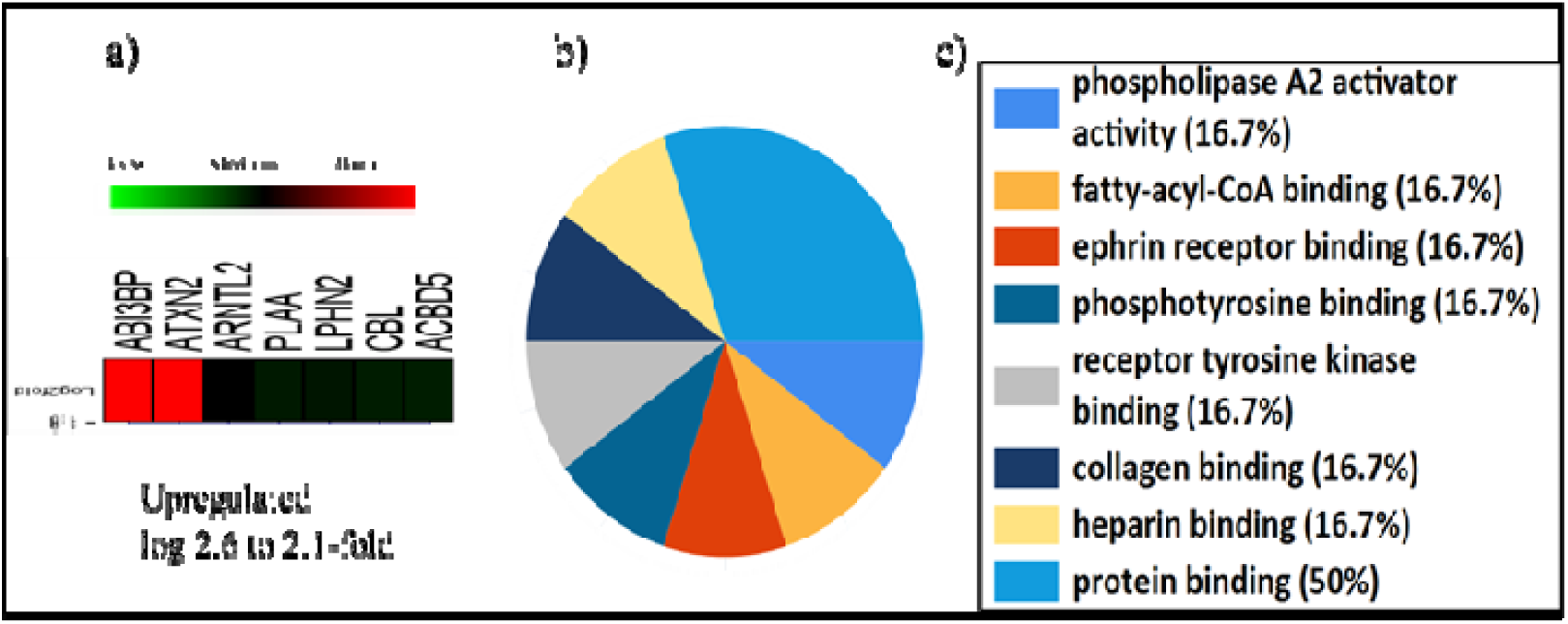
Differentially expressed genes from AmpliSeq transcriptome in Luminal B subtype a) Heatmap of DEGs, b) Pie charts showing the distribution of functions of DEGs, c) Different functions of the DEGs; included ECM component binding functions. Genes also showed the phospholipase A2 activator activity.

**Figure6 C:**
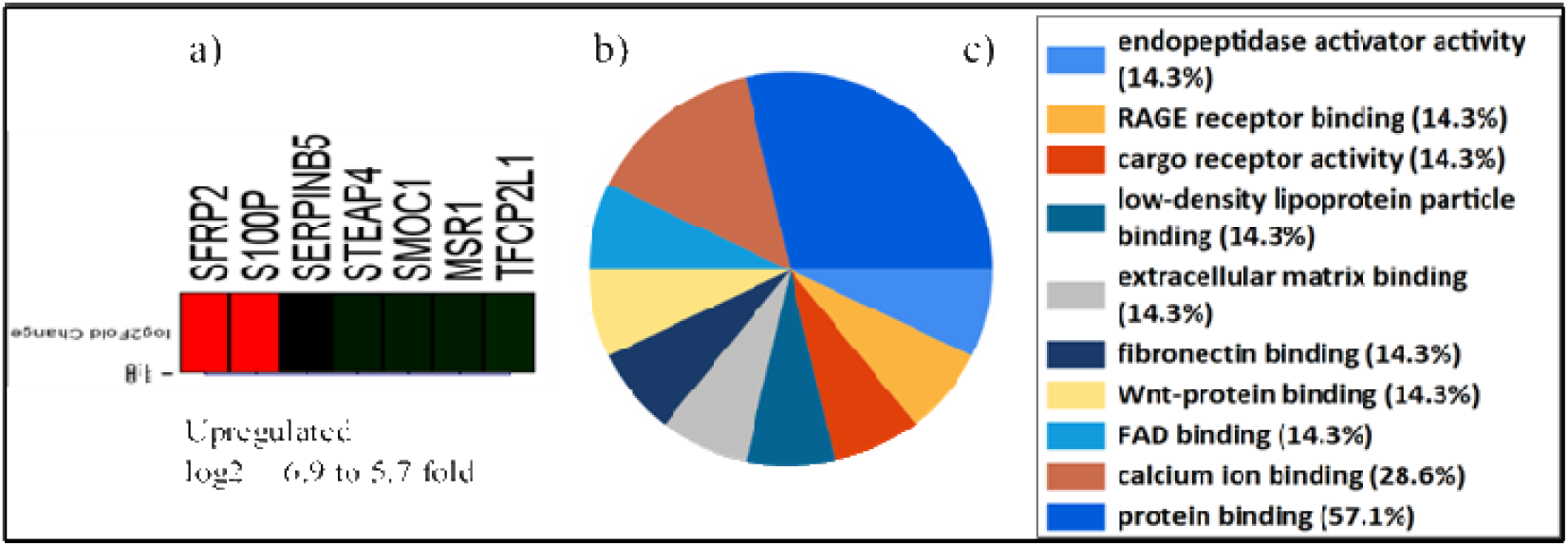
Differentially expressed genes from the AmpliSeq transcriptome in the Her-2 positive subtype. a) Heatmap of upregulated DEGs; b) Pie charts illustrating the distribution of functions of upregulated DEGs; c) Various functions of the upregulated DEGs, including G-protein receptor activity and leukotriene receptor activity. The genes also exhibit olfactory receptor activity and transcription factor activity; d) Pie charts depicting the distribution of functions of downregulated DEGs; e) Distinct functions of the downregulated DEGs include ECM structural components and collagen or ECM binding proteins

**Figure-6D:**
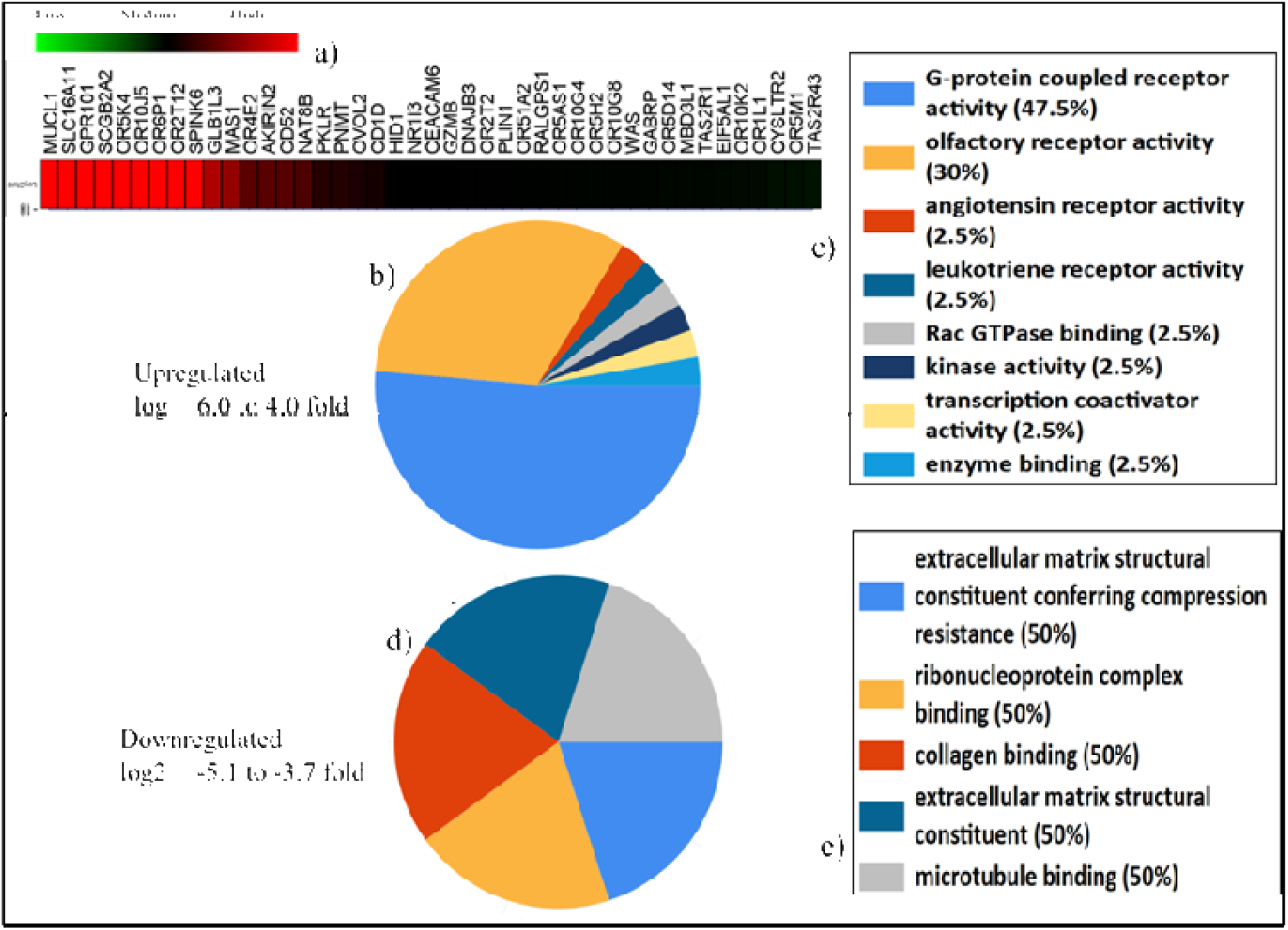
Differentially expressed genes from AmpliSeq transcriptome in TNBC subtype a) Heatmap of DEGs, b) pie charts showing the distribution of functions of DEGs, c) Different functions of the DEGs had fibronectin and calcium binding functions. Genes also showed endopeptidase activator activity.

A heatmap from both runs showed the CAF-derived genes with different molecular functions and their association with molecular subtypes. These findings suggest the existence of a distinct set of CAFs in TMEs across breast cancer subtypes and highlight their function heterogeneity.

### 3.2. Shortlisted genes and validation

Gene selection was conducted without bias. Genes exhibiting the highest expression fold change, robust statistical power (p-value & FDR ≥ 0.05), and significant functions in cancer were chosen for validation across additional patient cohorts. Based on their functions, these genes were categorized into three groups: (a) immune response-related genes: IL19, CD52, CD8A, MSR1, A2M, LILRB2, SLAMF-1; (b) ECM remodeling genes: MMP-3, MMP10, ASPM, FERMT1, MFAP5, COL11A1, GBRE, FN1, FBN1, DEGS2, CAPN9, SFRP2; and (c) calcium/ECM binding protein genes: CIB4, CTSK, CKAP5, S100P, ABI3BBP, ADAM33. The clinicopathological features of the patients enrolled for validation are provided in Table 2. Detailed biological functions of the shortlisted genes are presented in Supplementary Table 1.

**Table 2.** Clinicopathological characteristics of 83 cases used for validation

| Parameters | Luminal A<br>(<br>%<br>) | Luminal B (%) | Her-2+<br>(<br>%<br>) | TNBC (%) |
| --- | --- | --- | --- | --- |
| Number of cases | 25 | 18 | 18 | 22 |
| Age (years) |  |  |  |  |
| Mean | 53.75 ± 1<br>5<br>.<br>5<br>4<br>(Minimum<br>2<br>6<br>Maximum<br>8<br>4<br>) | 48 ± 15.1<br>(Minimum 21<br>Maximum 72) | 51 ± 11.78<br>(Minimum 3<br>4<br>Maximum 7<br>1<br>) | 49 ± 8.9<br>(Minimum 2<br>5<br>Maximum 6<br>2<br>) |
| Median | 54 ± 54.54 | 45 ± 15.1 | 53 ± 11.78 | 50 ± 8.9 |
| Histological type |  |  |  |  |
| IDC | 23 (92%) | 16 (89%) | 18 (100%) | 20 (90%) |
| ILC | 4% | 0 | 0 | 2 (10%) |
| Other | 4% (Solid | 2 (11%) | 0 | 0 |
| t | P | (Micr |  |  |
| y | a | opapi |  |  |
| p | p | llary, |  |  |
| e | il | Mixe |  |  |
| s | l | d |  |  |
|  | a | type) |  |  |
|  | r |  |  |  |
|  | y |  |  |  |
|  | C |  |  |  |

| Tumor size |  |  |  |  |
| --- | --- | --- | --- | --- |
| <2cm | 3 (12%) | 2 (11%) | 1 (5.5%) | 3 (14%) |
| 2-5cm | 17 (68%) | 12 (66.6%) | 15 (83.3%) | 16 (73%) |
| >5cm | 5 (20%) | 4 (22.2%) | 2 (11%) | 3 (14%) |
| Tumor grade |  |  |  |  |
| I | 3 (12%) | 0 (2.7%) | 0 | 0 |
| II | 16 (64%) | 6 (33.3%) | 2 (11%) | 5 (23%) |
| III | 6 (24%) | 12 (66.6%) | 16 (89%) | 17 (77%) |
| Nodal Status |  |  |  |  |
| Negative | 10 (40%) | 6 (33.3%) | 5 (28%) | 9 (41%) |
| Positive | 15 (60%) | 12 (66.6%) | 13 (72%) | 13 (59%) |
| Ki-67 (cut off 20%) |  |  |  |  |
| Low | 10 (40%) | 3 (16.6%) | 1 (5.5%) | 2 (10%) |
| High | 15 (60%) | 15 (83.3%) | 17 (94.5%) | 20 (90) |
| Follow-up (months) |  |  |  |  |
|  | 51.4 to 58 | 18 to 49.2 | 10 to 56 | 9.2 to 60.1 |
| Clinical stage |  |  |  |  |
| IA | 2 (16.6%) | 2 (22.2%) | - | - |
| IB | - | - | - | - |
| IIA | 4 (33.3%) | 3 (33.3%) | 4 (57%) | 6 (33.3%) |
| IIB | 1 (8%) | 2 (22.2%) | 2 (29%) | 10 (55.5%) |
| IIIA | 3 (25%) | 1 (11.1%) | 1 (14.2%) | - |
| IIIB | 1 (8.3%) | 1 (11.1%) | - | - |
| IIIC | 1 (8.3%) | - | - | 2 (11.1%) |
| Local recurrence |  |  |  |  |
|  | No | No | No | No |
| Distance recurrence |  |  |  |  |
|  | No | No | No | No |
| Menopause |  |  |  |  |
| Yes | 15 | 13 | 9 | 16 |
| No | - | - | - | - |
| Death |  |  |  |  |
|  | 1 | - | 1 | 1 |

#### 3.2.1. Immune response-related genes

Overall, we observed a varied mean expression fold change for upregulated genes (IL19, CD52, CD8A, MSR1), with a mean log from +2.3-fold to +11-fold; SD±1.1, and downregulated genes (A2M, LILRB2, SLAMF-1) with a mean log of -1.87-fold to -6.7-fold; SD±1.3. Notably, IL19, CD52, CD8A, and MSR1 exhibited high expression levels in the Luminal A & B and TNBC groups (up to mean log +11-fold), compared to the Her-2 positive group (mean log 3-fold) (Fig-7). Among these, IL-19, CD52, and MSR1 displayed the highest expression (log +8 to +11; SD±0.4) across both Luminal and TNBC subtypes. Conversely, LILRB2, SLAMF1, and A2M were downregulated in Luminal A/B and TNBC (mean log -6.7 to -4 fold; SD ± 0.5) compared to the Her-2 positive subtypes. When comparing within luminal groups, these genes showed slightly higher expression levels in luminal B than in luminal A. Based on this distinct pattern of CAFs expressing immune response-related genes, the breast cancer subtypes were categorized into two clusters: cluster A includes Luminal A, Luminal B, and TNBC, which comprised CAFs with high expression of immune response-related genes, while cluster B includes Her-2 positive, which contained CAFs with low expression of immune response-related genes. These findings highlight the unique distribution of CAFs with immune modulatory functions across the different breast cancer subtypes. Several prior studies have observed the tumor-promoting roles of the shortlisted immune response-related genes. For example, IL-19 fosters an immunosuppressive microenvironment in breast cancer by regulating the release of various pro-inflammatory cytokines and is linked to advanced tumor stages, increased metastasis, and poorer survival [22–23]. CD52 exhibited a positive correlation with CD8+ T cells, activated memory CD4+ T cells, macrophage M1, and Gamma Delta T cells, which may be crucial in sustaining the immune-dominant position of the tumor microenvironment [24].

Furthermore, H. Hu et al. (2021) identified a subset of CAFs that displayed high expression of CD52 [25]. Currently, elevated expression of CD52 was noted in luminal groups, which could be a factor contributing to better prognosis in luminal breast cancer patients compared to those with Her-2 positive and TNBC. A study identified CAFs expressing MSR1 and demonstrated macrophage phenotypes in breast cancer and glioma [26]. Consistent with previous research, SLAMF1 and A2M showed downregulated expression in this study [27–28]. In contrast to earlier findings, we observed downregulation of the LILRB2 gene. LILRB2 is a tumor-promoting gene, and its high expression correlates with proliferation and poor prognosis [29]. It may be that a different isoform of the LILRB2 gene, whose downregulation is linked to carcinogenesis, is involved in the current study Expression of immune response-related genes in CAFs reflects their immune cell-like phenotype and phenotypic plasticity characteristics. Phenotypic plasticity is one of the significant causes of therapy failure and relapse in aggressive cancer, including TNBC. The presence of such CAFs in the stroma of TNBC might be responsible for their aggressive feature and poor prognosis. Immune cell-like CAFs were also noted in the stroma of the luminal subtype, which possesses a slightly improved prognosis compared to TNBC. Such subtype-specific behavior of CAFs could be due to their selective cross-talk with epithelial cells or other stroma cells. Consequently, it would be interesting to see how and what distinct cytokines/chemokines milieu CAFs use to communicate with hormonal receptor positive and negative tumors.

#### 3.2.2. ECM remodeling genes

Overall, the mean expression fold change of upregulated ECM remodeling genes (MMP-3, MMP10, ASPM, FERMT1, MFAP5, COL11A1, GBRE, FN1, FBN1, SFRP2, FGFR2) ranged from log +4 to +11 fold (SD±1.1), while downregulated genes (DEGS2, CAPN9) ranged from log -2 to -7 fold (SD±1.2). Within subtypes, the comparison demonstrated that ECM remodeling genes exhibited significantly higher expression in the stroma of Her-2 positive and TNBC subtypes (log +4 to +11-fold; SD±0.6) compared to Luminal subtypes (log +1.5 to +6-fold) (Fig. 8). Notably, COL11A1 showed the highest expression (log +11-fold; SD ± 1.2) in both Her-2 positive and TNBC subtypes. COL11A1, a component of type XI collagen and a major ECM element, is secreted by CAFs. Previous studies have frequently reported high COL11A1 levels in various cancers correlated with poor clinical outcomes [30]. Interestingly, COL11A1 can influence immune cell infiltration in the TME and is often associated with immunosuppressive traits, chemoresistance, and recurrence [30]. Consequently, elevated COL11A1 levels may contribute to poor prognosis in Her-2 positive and TNBC due to an immunosuppressive TME. Other overexpressed ECM remodeling genes have been documented for their potential in metastatic initiation, EMT activation, angiogenesis promotion, immune infiltration, and immunosuppressive functions. All these cancer hallmarks are well characterized in the highly aggressive subtypes of TNBC and Her-2 positive breast cancer. Thus, the expression of these signatures in the stroma of TNBC and Her-2 positive emphasizes the tumor-promoting role of CAFs

**Figure 7:**
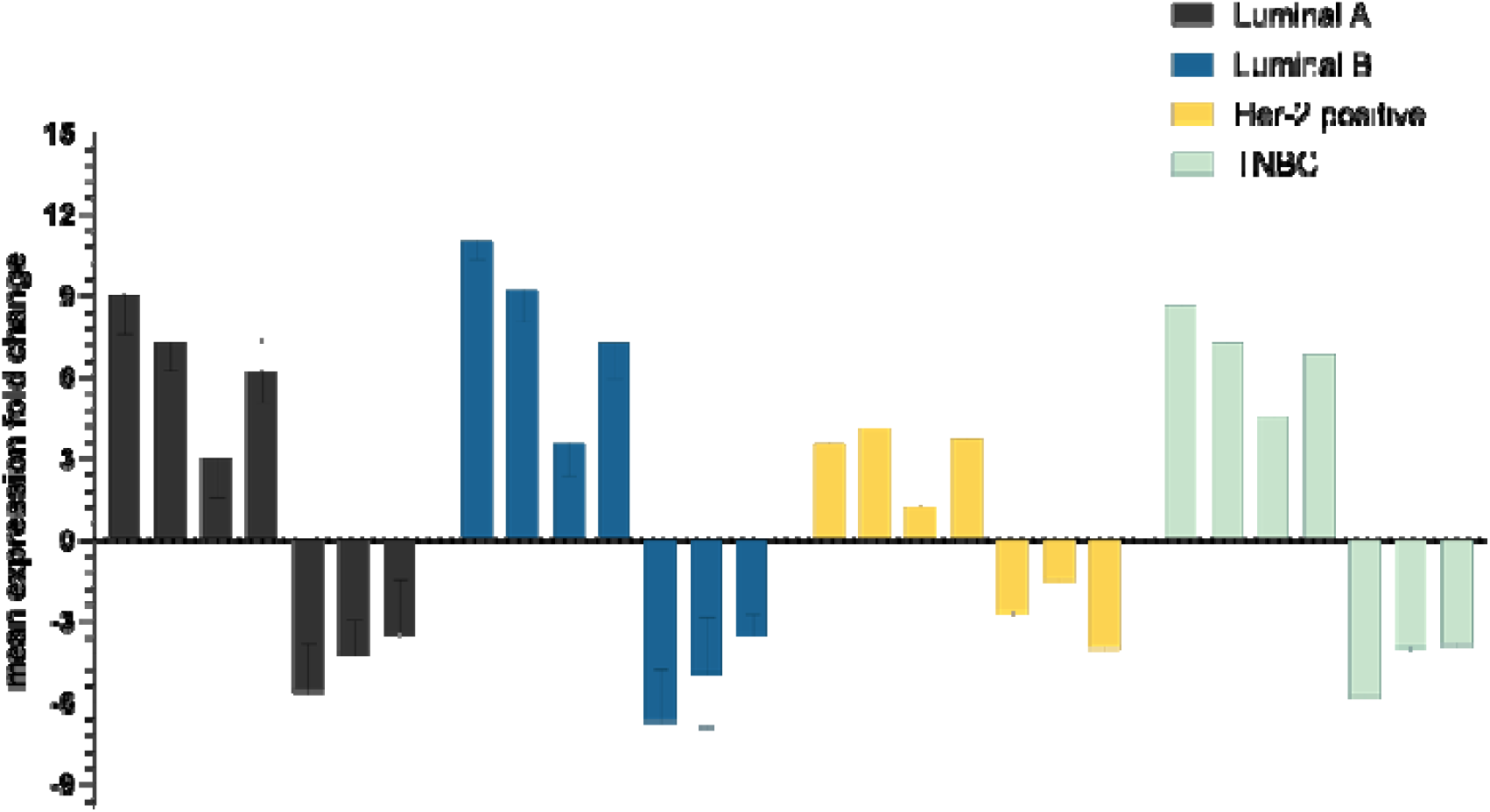
Shows the mean fold change expression of immune response-related genes across the breast cancer subtypes.

**Figure 8:**
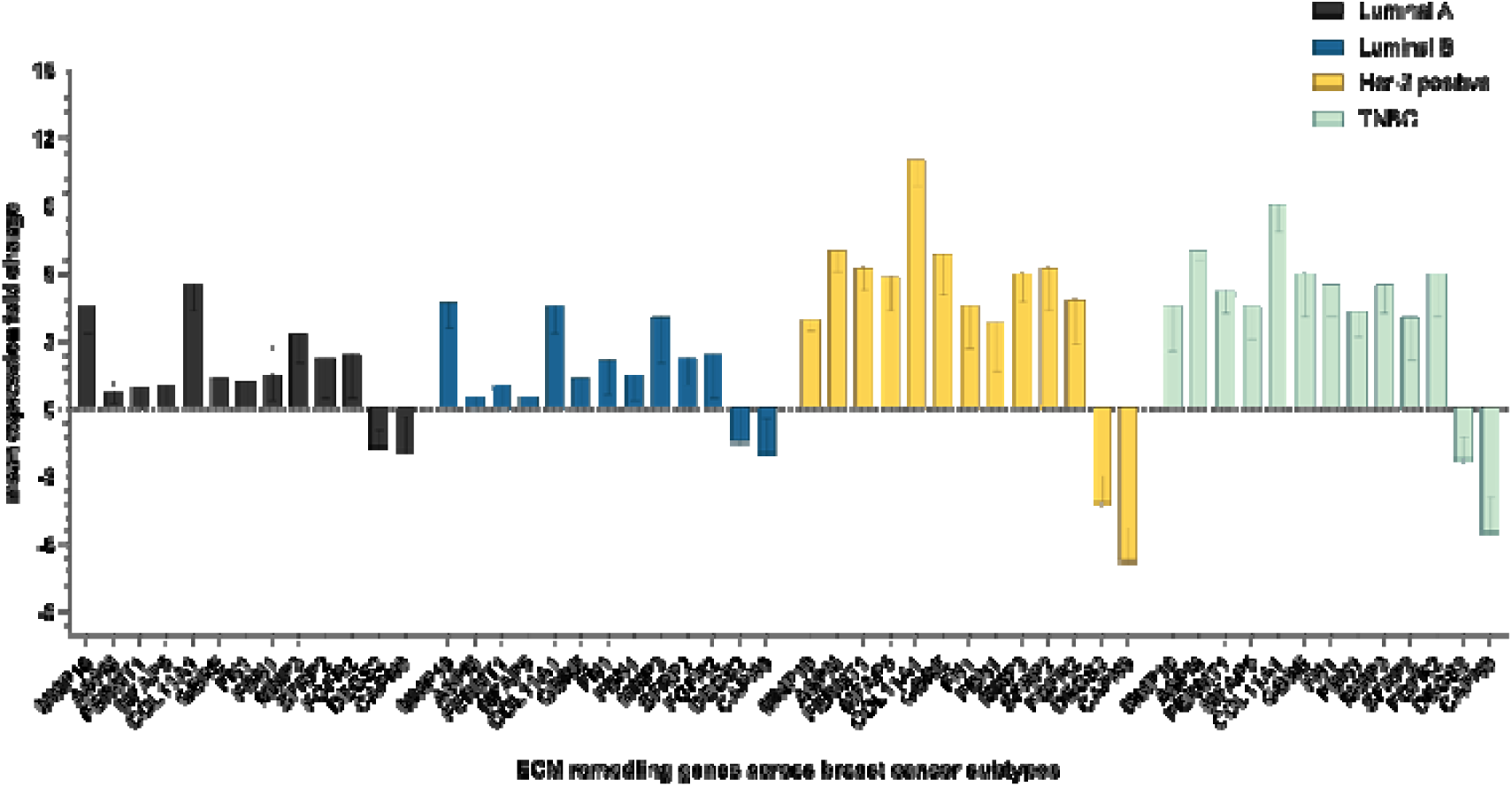
Shows the mean fold change expression of ECM remodeling genes across the breast cancer subtypes.

For instance, MMPs (MMP-3 and MMP-10), FERMT1, MFAP5, and GABRE are linked to a high metastatic potential [31–37]. GABRE can also enhance the release of MMPs in the extracellular matrix (ECM) and thus directly contribute to the metastatic activation role of cancer-associated fibroblasts (CAFs) [36–37]. In pancreatic cancer, MFAP5 has been shown to modify the effects of PD-L1 therapy [35]. FN1 demonstrated the ability to induce immune cell infiltration in the tumor stroma [38–39]. These findings suggest that MFAP5 and FN1 may function together to support “immune response-related genes.” Additionally, MFAP5 and FERMT1 play a role in epithelial-mesenchymal transition (EMT) activation and could contribute to therapy resistance in Her-2 positive and triple-negative breast cancer (TNBC). Similarly, FBN1 and SFRP2 have been identified as activators of angiogenesis in cancer [40–41]. Our prior study reported elevated microvessel density (MVD) in the same Her-2 positive and TNBC patients [42]. Moreover, FGFR2 can activate several downstream pathways, including RAS–MAPK and PI3K/AKT, thereby contributing to a high proliferation rate in Her-2 positive and TNBC [43–44]. On the other hand, CAPN9 and DEGS2 were relatively downregulated in Her-2 positive and TNBC compared to luminal subtypes. CAPN9, a tumor suppressor gene, has previously been linked to poor prognosis in hormone receptor-positive breast cancers due to its low expression [45]

Based on the differential expression of CAF-derived ECM remodeling genes, breast cancer subtypes were divided into two clusters: Cluster A includes Her-2 positive and TNBC subtypes with high ECM remodeling gene expression, while Cluster B consists of Luminal A and Luminal B subtypes with low ECM remodeling gene expression. Thus, the underlying ECM remodeling functions of CAFs have a unique correlation with subtypes.

#### 3.2.3. Calcium/protein binding genes

Calcium-binding genes exhibited generally similar expression across various breast cancer subtypes (Fig. 9). Upregulated genes (CIB4, CTSK4, S100P, and AB13BP) demonstrated mean expression fold changes of log +4 to +7-fold; SD ± 1.4. Conversely, downregulated genes (ADAM33 and CKAP5A) indicated mean expression fold changes ranging from log -5.5 to -3.5-fold; SD ± 1.3. Thus, a subset of CAFs expressing adhesion molecules was evenly distributed across the breast cancer subtypes. Previous studies have identified the pro-tumorigenic role of this category of genes. S100P enhances the aggressive properties of breast cancer cells in Her-2 positive breast cancer patients [46]. High mRNA expression of S100P has been linked to the poorest overall survival (HR = 1.46, 95% CI: 1.15–1.85, p = 0.0017) [47]. CIB4 encodes another calcium-binding protein that regulates Ca2+ ion efflux or their mediated signaling pathways, like cellular proliferation, migration, and angiogenesis. This leads to chemotherapy resistance, predominantly associated with TNBCs and Her-2 positive breast cancer—an expression of CTSK4 and ASPM in CAFs, which indicates a high rate of cell division in CAFs. Given that cancer cells typically divide within the tumor mass, it is possible that CTSK4 and ASPM-expressing CAFs originated from tumor cells in proximity to epithelial cells [48]. This also underscores the presence of another CAF subpopulation characterized by high cell cycle division. ABI3BP is an ECM-binding protein [49]. Yan F et al. (2022) investigated the dysregulated expression of ABI3BP in lung carcinoma, which was closely related to immune cell infiltration and suggested it might predict the therapeutic effect of immune checkpoint inhibitors [50]. Thus, CAFs expressing the ECM-binding protein ABI3BP also exert an immune modulatory effect. Moreover, ADAM33 and CKAP5A (−5.5 to -3.5-fold; SD ± 1.3) were downregulated across the selected validation cases. Previously, Manica G et al. (2017), through in vitro and in vivo studies, associated the lower expression of ADAM33 in TNBC and BLBC patients with shorter overall survival [51]. Lan J et al. (2023) also observed a downregulation of ADAM33 in 139 thyroid cancer biopsy samples [52]. Furthermore, CKAP5A (log -3.2; SD±1.2 in Luminal A to -4.2-fold; SD ±1.01 in TNBC) is an essential microtubule-associated protein, and its silencing leads to chemotherapy resistance in an ovarian cancer cell line due to a reduction in EB1 dynamics during mitosis [53]. Schneider et al. (2017) also identified the dysregulation of CKAP5 and its association with the prognosis of NSCLC patients [58].

**Figure 10:**
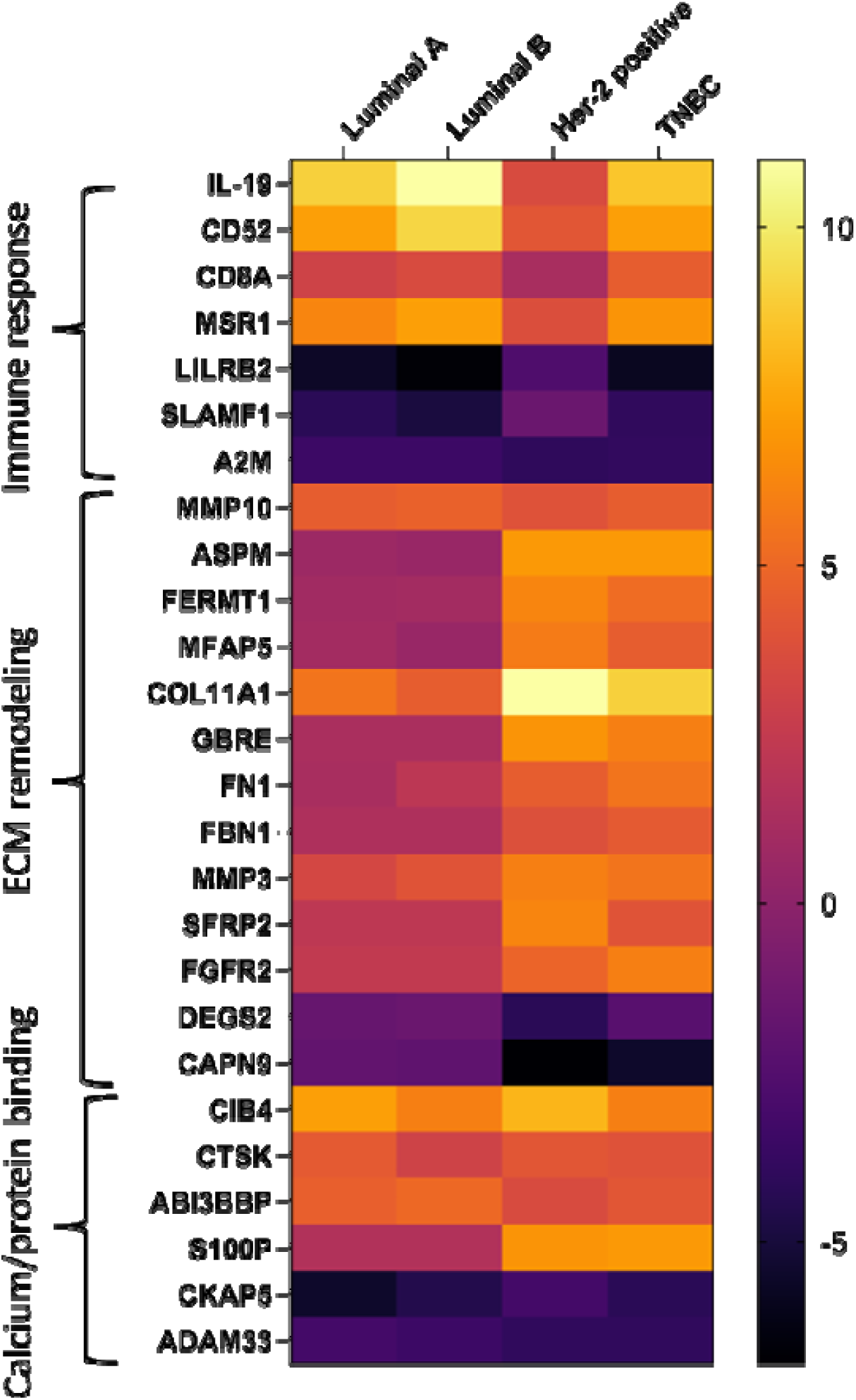
Combined heatmap of ECM remodeling, immune response, and calcium binding genes, showing the differentially expressed genes in intrinsic breast cancer subtypes.

When these gene signatures were correlated with clinicopathological parameters, no statistically significant correlations were found with tumor size, lymph node status, histological grade, or histology subtype.

### 3.3. Protein-protein interaction network

The biological interactions among selected genes from the three categories at the protein level were also examined. The online tool STRING was utilized to observe the protein-protein interaction network. Genes from the ECM remodeling and immune response groups demonstrated specific interactions at the protein level. The ECM remodeling genes MFAP5, FBN1, COL11A1, MMP3, and MMP10 exhibited biological interactions at the protein level. According to the gene ontology database, these interactions make up the extracellular matrix structural remodeling (collagen or ECM degradation) pathways. This suggests their important role in tumor cell invasion and metastasis. The immune response genes MSR1, CD52, CD8A, LILRB2, and CD79A showed biological interaction at protein levels (Supplementary-2). According to the KEGG pathways and UniPort databases, these interactions constituted signaling pathways with adaptive immunity functions.

## 4. Discussion

The tumor microenvironment (TME) is recognized as a vital component of the tumor mass. It comprises stromal cells such as cancer-associated fibroblasts (CAFs), immune cells, tumor cells, adipocytes, and endothelial cells, along with the various cytokines, growth factors, hormones, transcription factors, and proteases they secrete. The TME holds therapeutic and diagnostic potential. Among all cell types within the TME, CAFs are critical due to their significant roles in extracellular matrix (ECM) remodeling, immune cell infiltration, immunosuppression, metastasis, and angiogenesis. CAFs are the most heterogeneous TME cells stemming from multiple cell origins and phenotypic functions, which complicates targeted therapies. This CAF heterogeneity is particularly pronounced in breast cancer, which is classified into four intrinsic molecular subtypes based on gene expression profiling. We hypothesized that, like tumor cells, CAFs exhibit molecular heterogeneity, different intrinsic subtypes of breast cancer maintaining distinct clusters of CAFs that express stromal genes. In this study, we evaluated the whole transcriptome of tumor tissue to emphasize the differential expression of stromal genes across molecular subtypes.

Additionally, we assessed the AmpliSeq transcriptome of pure CAFs isolated from primary human breast tumors of various molecular subtypes. In both whole transcriptome and AmpliSeq analyses, we identified breast cancer subtype-specific CAF populations based on their functional phenotypes. Genes associated with CAFs showed enrichment in diverse molecular functions and demonstrated differential expression across breast cancer subtypes. Consequently, based on functional gene enrichment analysis, these CAFs were categorized into three subpopulations: those related to immune response, ECM remodeling, and calcium/protein binding. Our findings highlight the existence of three functionally distinct subpopulations of CAFs in breast cancer: first, those that modulate the immune response and assist cancer cells in evading immune surveillance; second, those involved in ECM synthesis and remodeling that facilitate invasion and metastasis; and third, those that regulate cancer signaling pathways by managing Ca2+ ions and binding to other proteins (ECM proteins and proteases)

Previously, Ӧhlund and colleagues [19] discovered and identified two distinct subsets of cancer-associated fibroblasts (CAFs) in pancreatic cancer: myofibroblastic CAFs (myCAFs) and inflammatory CAFs (iCAFs). Researchers also identified another CAF subpopulation, “antigen-presenting CAFs” (apCAFs), which exhibited MHC class II and CD74 expression instead of traditional costimulatory molecules. CAF subpopulations have also been reported in breast cancer. For instance, four different CAF subsets (S1-S4) are classified based on their varying expression of fibroblast markers, such as CD29, FAP, α-SMA, PDGFRβ, FSP1, and CAV1. Bartoschek et al. [21] defined four more CAF subpopulations: vCAFs, mCAFs, cCAFs, and dCAFs, according to their distinct cellular sources identified through single-cell RNA sequencing. Similar to previously published data, the current study also identified at least two CAF subpopulations, namely, immune modulatory and ECM remodeling

Furthermore, we have validated the functionally important short-listed genes from both runs using a real-time PCR approach in 83 breast cancer samples. In validation, immune response-related genes were highly enriched in TNBC and Luminal subtypes compared to the Her-2 positive subtype; ECM remodeling genes were over-expressed in TNBC, while Her-2 positive and Calcium/protein binding genes exhibited overall similar expression across the molecular subtypes. These findings highlighted the nonuniform distribution of CAFs across the breast cancer subtypes, thus supporting the hypothesis that CAF functional heterogeneity exists in the breast cancer microenvironment. The validation experiment showed the expression of CAF-derived essential tumor promoter genes in highly aggressive breast cancer subtypes, such as Her-2 positive and TNBC. Moreover, the genes chosen for validation have demonstrated pro-tumorigenic roles. For instance, IL-19, part of the IL-10 family of cytokines, can induce an immunosuppressive microenvironment in breast cancer, and its elevated expression has been linked to advanced tumor stages, increased metastasis, and poor survival. Studies by CH Hsing et al. (2012) and colleagues indicated that IL-19 induced the transcription of IL-1β, IL-6, TGF-β, MMP-2, MMP-9, and CXCR4 genes in breast and esophageal cancer cells, as well as fibronectin expression and assembly [22–23]. CD52, expressed on the surface of many lymphoid and other hematopoietic cells, influences the prognosis of breast cancer patients [24]. A soluble form of CD52 induces immunosuppression by inhibiting the activation and proliferation of immune cells, particularly CD4+ T cells, toll-like receptor, and TNF-α receptor pathways, as well as the production of NF-kB and pro-inflammatory factors [24]. Consistent with the present study, another investigation identified a subpopulation of CAFs with high CD52 expression in lung cancer [25]. Additionally, MSR1 (CD204) plays crucial roles in inflammation modulation across multiple pathophysiological processes, including M2 polarization [26]. MSR1 serves as a marker for M2 pro-inflammatory macrophages, and its expression on CAFs indicates a macrophage-like role of CAFs in breast cancer [26]. In glioma, MSR1 positive CAFs were associated with a higher risk and worse prognosis [54]. Similarly, LILRB2, which exhibited lower expression in the present study, interacts with HLA class I molecules and correlates with immune and monocytic lineage gene signatures. Upregulation of LILRB2 in macrophages offers an evasion mechanism for cancer cells against phagocytosis [29]. Previous studies reported that LILRB2 functions as an immune inhibitory receptor, activates MDSCs, and promotes M2-like phenotypes. Furthermore, SLAMF1, a signaling lymphocytic activation molecule 1, had decreased expression in patients with aggressive chronic lymphocytic leukemia (CLL) and was associated with reduced overall survival [27]. Patients with high SLAMF1 expression in colorectal cancer had a significantly higher survival rate than those with low expression, suggesting that SLAMF1 is an anti-tumor biomarker in CRC [55]. A2M has a well-documented role as a protease inhibitor, immune modulator, and scavenger of growth factors, hormones, and cytokines [28]. Therefore, CAFs with upregulated genes IL-19, CD52, CD8A, and MSR1, along with the downregulation of genes LILRB2, SLAMF1, and A2M, as observed in this study, may induce an immunosuppressive microenvironment in TNBC and Luminal subtypes. Unfortunately, we could not correlate these CAF signatures with the clinicopathological features of the patients. However, Finak et al. developed a 26-gene predictor (stroma-derived prognostic predictor) that offers more accurate predictions of disease outcomes [56]. In their study, the tumor stroma from the good outcome cluster overexpressed a distinct set of immune-related genes, including T cell and NK cell markers, indicative of a Th1–type immune response (Granzyme A (GZMA), CD52, CD247, CD8A). In contrast, the poor prognosis group exhibited high expression of hypoxia and angiogenesis genes alongside low expression of type I immune response genes [56]. This finding reinforces the clinical significance of CAF signatures observed in the present study. It suggests that individuals with this gene expression pattern might benefit from treatments targeting tumor cells via the immune response, such as vaccine therapies and immunotherapy.

Moreover, ECM remodeling genes (MMP-3, MMP10, ASPM, FERMT1, MFAP5, COL11A1, GBRE, FN1, FBN1, S100P, DEGS2, CAPN9, SFRP2, and COL11A1), which exhibited high expression in Her-2 positive and TNBC, have pro-tumorigenic roles through the activation and regulation of invasion, migration, metastasis, angiogenesis, EMT, and immune cell infiltration. Previous studies have noted that COL11A1 is linked to advanced cancer stages and can regulate immune cell infiltration in breast cancer [30]. Similarly, overexpressed MMP-3/10, FERMT1, MFAP5, and GABRE may promote metastasis, angiogenesis, EMT, and immune response in both Her-2 positive and TNBC. MMP3/10 catalyzes the degradation of various ECM substrates, including proteoglycans, laminin, fibronectin, and collagen IV, facilitating angiogenesis and metastasis [31]. L Li et al. (2022) confirmed that FERMT1 can regulate EMT via NLRP3 and inhibit the NF-kB signaling pathway [32]. Studies indicate that MFAP5 is crucial for integrating elastic microfibers and regulating endothelial cells. MFAP5 can promote EMT in BLBC metastasis through the TGF-β/Notch pathway [33–34]. Y Duan et al. (2023) even demonstrated that targeting MFAP5 high CAFs might serve as a potential adjuvant therapy to enhance the immunochemotherapy effect in PDAC by remodeling the desmoplastic and immunosuppressive microenvironment [35]. Finally, a study found that increased expression of GABRE is implicated in cancer metastasis by promoting MMP production in cancer cells [36–37]. Likewise, FN1, a glycoprotein found throughout the ECM, plays a critical role in the initiation and progression of cancer and has been associated with immune cell infiltration and unfavorable responses to immune checkpoint inhibitors in breast cancer treatment [38–39]. Similarly, FBN1 has been identified in a novel TGF-β-induced marker and knockdown study by HC Lien et al. (2019), which showed a reduction in migration, invasion, angiogenesis, and EMT [40]. Angiogenesis highlights the role of SFRP2 in breast cancer, as noted by another study [41]. Furthermore, FGFR2 is a membrane-spanning tyrosine kinase that plays a crucial role in breast tumorigenesis by modulating primary downstream cancer signaling pathways such as RAS–MAPK and PIK3CA/AKT [43–4]. Moreover, CAPN9 and DEGS2 were the downregulated genes identified in the current study; CAPN9 had slightly higher expression in Luminal A/B than in Her-2 positive and TNBC. In alignment with our research, J Davis et al. (2014) determined its low expression and additionally implicated it in poor prognosis in HR+ breast cancer after endocrine therapy [45]. Perhaps, under current conditions, the basal expression of CAPN9 in Luminal cases could account for a favorable prognosis in HR+ cases compared to HR-. This is supported by the work of Peng P., who in 2016, noted low expression levels of CAPN-9 in gastric cancer patients with unfavorable prognoses [57].

Following validation, the calcium/protein binding genes in the CAFs category revealed that S100P, CIB4, CTSK4, and AB13BP were upregulated, while ADAM33 and CKAP5A were uniformly downregulated across breast cancer subtypes. All these genes are well-characterized for their significant roles in cancer progression. S100P, a calcium-binding protein, promotes an aggressive phenotype in Her-2 positive breast cancer patients and is linked to their worst overall survival [46–47]. CIB4 can also regulate Ca2+ ion efflux-mediated cancer signaling pathways. Furthermore, CTSK4 encodes the CDK4 enzyme, which, along with ASPM, promotes the proliferation of CAFs in the breast tumor microenvironment (TME) and enhances the pro-tumorigenic functions of CAFs [48]. Moreover, aberrant expression of ABI3BP has been correlated with immune infiltration, response to immune checkpoint inhibitors, and prognosis in renal chromophobe carcinoma, mesothelioma, pancreatic adenocarcinoma, and lung cancer [49–50]. Thus, we conclude that a subset of CAFs with “calcium/protein binding function” also operate functionally with a subgroup of CAFs related to “immune response.” Additionally, ADAM33 and CKAP5A were observed to be downregulated in the validation cohort. Previous studies have correlated the downregulation of ADAM33 with shorter overall survival in breast cancer (TNBC and BLBC) and thyroid cancer patients [51–52]. Although ADAM33 is an oncogene, its downregulation does not contribute to pro-tumorigenic functions. It may involve a different isoform of ADAM33 whose downregulation induces tumorigenic functions [52]. Furthermore, Schneider MA et al. (2017) analyzed the downregulation of CKAP5 in lung cancer cases, noting that five specific mitosis-associated genes (AURKA, DLGAP5, TPX2, KIF11, and CKAP5) correlate with poor prognosis for non-small cell lung cancer patients. CKAP5 is a microtubule-associated protein, and its knockout-caused chemo-resistance in ovarian cancer cell lines [53]. Thus, it is currently found that a subpopulation of CAFs contributes to breast cancer carcinogenesis through the aberrant expression of calcium/protein binding genes, regardless of molecular subtypes.

Furthermore, STRING analysis revealed that ECM remodeling genes MFAP5, FBN1, COL11A1, MMP3, and MMP10 interact biologically, potentially forming the ECM remodeling pathways (KEGG pathways and UniPort data) involved in tumor cell invasion and metastasis. Likewise, immune response genes MSR1, CD52, CD8A, LILRB2, and CD79A also exhibited biological interactions at the protein level, forming signaling pathways related to adaptive immune functions. Although the current study employed high-throughput RNASeq methods and identified different functional populations of CAFs and their distinct connections to breast cancer, it showcases the general phenotypes of CAFs. Utilizing a newly developed single-cell sequencing method combined with spatial transcriptomics can provide deeper insights into the heterogeneity of CAFs in the TME. It can elucidate their mode of cross-talk with other cells.

## 5. Conclusion

This study identified three significant populations of CAFs (immune response-related, ECM remodeling, calcium-binding), highlighting the heterogeneity of CAFs in breast cancer. After validating the shortlisted genes from the CAF populations mentioned above, an immune response-related gene signature was observed in Luminal A/B and TNBC subtypes. The ECM remodeling gene signature was noted in Her-2 positive and TNBC subtypes. A separate subset of CAFs in the “calcium-binding population” was also identified, exhibiting high cell division capacity. The protein-protein interaction network suggested the roles of the shortlisted ECM remodeling genes in tumor cell invasion and metastasis and the immune response-related genes in signaling pathways associated with adaptive immunity functions. Thus, this study proposes that, based on the specific CAF population of each subtype, molecularly targeted therapy is necessary to address the stromal components that enhance tumor progression.

## Institutional Review Board Statement

The Institutional Ethical Committee (IEC) approved the study vide No. NK/4116/PhD/927, dated 31/12/2021; PGIMER, Chandigarh-160012, INDIA. Informed consent was obtained from each patient enrolled in the study.

## Informed Consent Statement

An informed consent was obtained from all subjects involved in the study.

## Data Availability Statement

The data presented in the study was generated by the author and not taken from other resources. The author will make the data available upon request.

## Acknowledgments

The author would like to acknowledge ICMR, New Delhi, India, and PGIMER, Chandigarh, India, for their support in completing this study.

