## Supplementary material for "Deciphering Functional Heterogeneity of Cancer-Associated Fibroblasts Across Molecular Subtypes of Breast Cancer": Supplemantory: Supplementary file_15.05.2024.docx

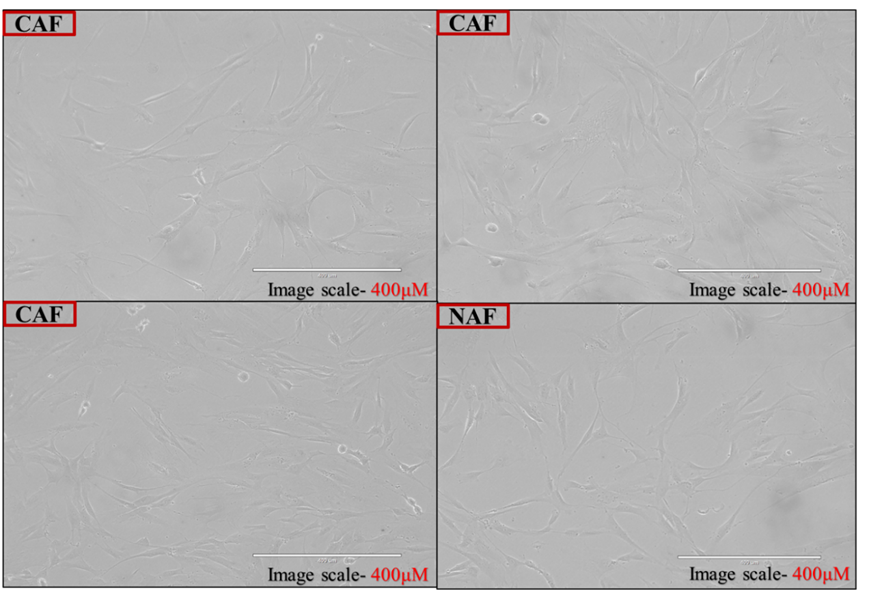

***Supplementary Figure:* Images of primary fibroblast culture isolated from breast cancer and normal breast tissue. CAF (Cancer-associated fibroblast), NAF (normal adjacent fibroblasts, isolated from normal adjoining breast tissue)**

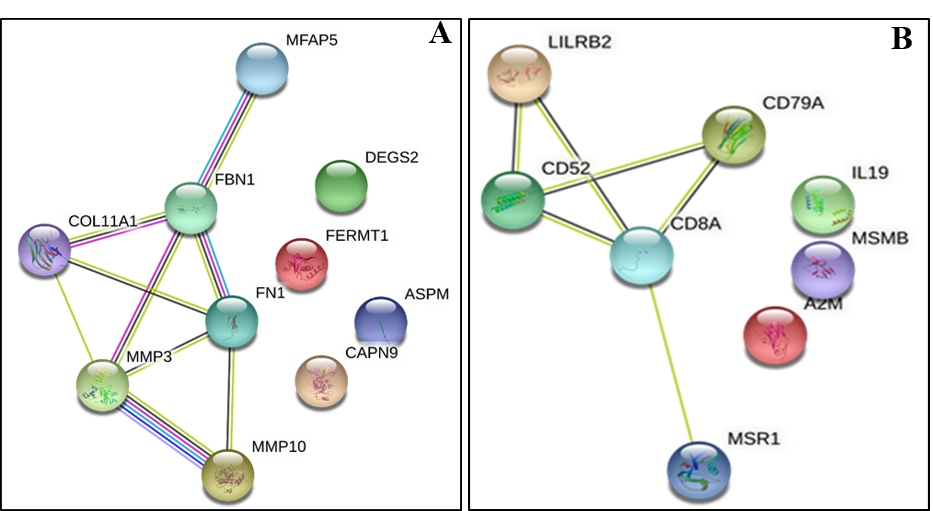

**Figure 2: A: Image of protein-protein interaction network of ECM remodelling genes generated form with help of STRING online tool. The MFAP5, FBN1, COL11A1, MMP3, and MMP10 ECM remodelling genes had biological interaction at protein levels. B: Protein-protein interactions network of immune response genes generated from with help of STRING online tool. The LILRB2, CD52, CD8A, CD79A, and MSR1 immune response genes had biological interaction at protein levels.**

| **Supplementary Table 1- List of the selected genes with their biological function** | | | | |
| --- | --- | --- | --- | --- |
|  | **Upregulated** | | **Downregulated** | |
| **Luminal A** | **Gene Name** | **Function** | **Gene Name** | **Function** |
|  | CIB4 | Calcium binding | A2M | Immune response |
|  | CTSK | Collagenase binding |  |  |
|  | MMP-3 | Matrix metalloproteinase |  |  |
| **Luminal B** | IL19 | Immune response |  |  |
|  | ABI3BBP | Collagen & Heparin-binding |  |  |
|  | CD52 | Immune response |  |  |
| **Her-2 positive** | MMP10 | Matrix Metalloproteinase | DEGS2 | Sphingosine hydroxylase activity |
|  | ASPM | ECM remodelling | CAPN9 | Sphingosine hydroxylase activity |
|  | FERMT1 | ECM remodelling | CKAP5 | Micro tubulin binding |
|  | MFAP5 | ECM remodelling |  |  |
|  | COL11A1 | ECM remodelling |  |  |
|  | GBRE | ECM remodelling |  |  |
|  | FN1 | ECM remodelling |  |  |
|  | FBN1 | ECM remodelling |  |  |
| **TNBC** | SFRP2 | Fibronectin binding |  |  |
|  | S100P | Fibronectin binding | LILRB2 | Immune response |
|  | MSR1 | Calcium binding | SLAMF-1 | T-cell receptor |
|  | CD8A | Immune response | ADAM33 | MMP-28 |
